# Long-term implicit memory for sequential auditory patterns in humans

**DOI:** 10.1101/2020.02.14.949404

**Authors:** Roberta Bianco, Peter M. C. Harrison, Mingyue Hu, Cora Bolger, Samantha Picken, Marcus T. Pearce, Maria Chait

**Author notes:** Correspondence concerning this article should be addressed to Roberta Bianco, UCL Ear Institute, University College London, 332 Grays Inn Rd, London, UK, WC1X 8EE. Author Note: Peter Harrison is now at the Max Planck for Empirical Aesthetics, Frankfurt, Germany.

## Abstract

To understand auditory scenes, listeners track and retain the statistics of sensory inputs as they unfold over time. We combined behavioural manipulation and modelling to investigate how sequence statistics are encoded into long-term memory and used to interpret incoming sensory signals. In a series of experiments, participants detected the emergence of regularly repeating patterns in novel rapid sound sequences. Unbeknownst to them, a few regular patterns reoccurred sparsely (every ∼3 minutes). Reoccurring sequences showed a rapidly growing detection time advantage over novel sequences. This effect was implicit, robust to interference, and persisted up to 7 weeks. Human performance was reproduced by a memory-constrained probabilistic model, where sequences are stored as n-grams and are subject to memory decay. Results suggest that similar psychological mechanisms may underlie integration processes over different-time scales in memory formation and flexible retrieval.

## Introduction

Memory is a crucial component of sensory processing, on multiple processing levels (Bale et al., 2017; Muckli & Petro, 2017). In the auditory modality, the ability to identify essentially any sound source, from footsteps to musical melody, requires the capacity to hold consecutive events in memory so as to link past and incoming information into a coherent emerging representation (Koelsch, Vuust, & Friston, 2019; McDermott, Schemitsch, & Simoncelli, 2013; Winkler, Denham, & Nelken, 2009). Whilst traditional models of sensory memory (e.g. Cowan, 1998) argued that such sensory traces are characterized by short retention times and computational encapsulation, a large body of work has since revealed that observers can retain detailed sensory information implicitly, over long periods (Arciuli & Simpson, 2012; Chun, 2000; Jiang, Song, & Rigas, 2005; Kim, Seitz, Feenstra, & Shams, 2009; Lu & Vicario, 2014; Winkler & Cowan, 2005). A compelling instance of this was demonstrated by Agus et al. (2010; see also Agus & Pressnitzer, 2013; Kang, Agus, & Pressnitzer, 2017) who showed that naïve listeners readily remembered certain spectro-temporal features of random noise bursts, such that reoccurring snippets were recognized weeks after initial exposure.

Here, we focus on long-term memory formation for arbitrary frequency patterns within rapidly unfolding sequences of discrete sounds. We ask whether naïve listeners can become sensitized to sparsely reoccurring tone sequences and investigate the conditions under which such memories are formed. To formalise the underlying psychological mechanisms, we compared human performance with a probabilistic model of sequential memory (Harrison, Bianco, Chait, & Pearce, 2020; Pearce, 2018).

The experimental design (Fig. 1) capitalizes on a paradigm developed by Barascud et al (2016) for measuring listeners’ sensitivity to complex acoustic patterns. Using fast sequences of short tones, they showed that listeners can rapidly detect the transition to a regularly repeating pattern (REG) from a sequence of random tones (RAN). Sequences were all novel and too rapid to allow for conscious tracking, but on most trials, participants were able to respond soon after the onset of the second cycle of regularity, implicating an efficient memory for the immediate sequence context. Here we ask how this memory is affected if the tone pattern was already experienced in the past.

**Fig. 1.**
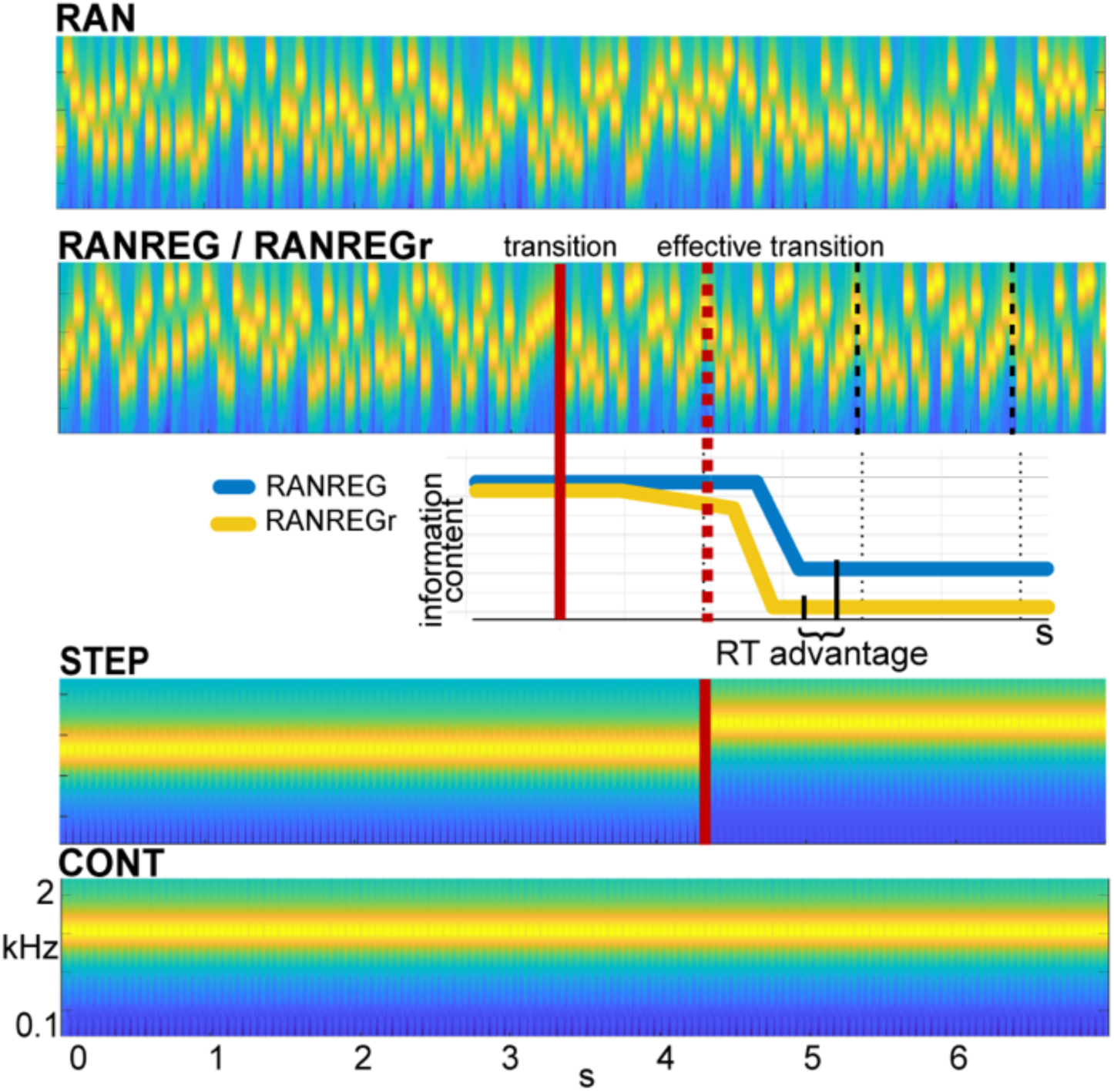
Example stimuli. Sequences were generated anew on each trial from a pool of 20 tone-pips of 50 ms duration each. RAN sequences were generated by randomly sampling from the full pool with replacement; RANREG sequences contained a transition from a random (RAN) to a regularly repeating cycles of 20 tone-pips (REG, cycles are marked with dashed lines). Therefore, the transition was manifested as a change in pattern only whilst maintaining the same long-term first-order statistics. The transition (randomized between 3 – 4 s post onset) is indicated by a red line; the red dashed line marks the ‘effective’ transition – the point at which the pattern starts repeating and hence becomes statistically detectable. Participants were instructed to respond to such transitions (50% of trials) as soon as possible. STEP stimuli, containing a step change in frequency (and their ‘no change’ control, CONT) were also included in the stimulus set for the purpose of estimating simple reaction time. Three (six in Exp. 4 and Exp. S1) particular regular patterns (REGr) were presented identically across 3 trials within a block (RANREGr). Reoccurrences were spaced ∼2.7 min apart. Different REGr were used for each participant. A schematic representation of outputs from the ideal observer (IDyOM) model is provided to illustrate how pattern reoccurrence might affect reaction time. For each tone in a sequence, the model outputs information content (IC) as a measure of its unexpectedness, given the preceding context. After the transition from a RAN to REG pattern, the IC drops over a few consecutive tones, reflecting the discovery of the REG. The brain is hypothesized to be sensitive to this change in IC, and once sufficient evidence has been accumulated, the emergent regularity ‘pops out’ perceptually. Therefore, RTs to onset of regularities can be used to quantify the amount of sensory information (number of tone-pips), required to detect the increasing predictability within the unfolding sequence. The black solid lines indicate the crossing of this putative evidence threshold. For novel patterns (blue line) this typically occurs within the first cycle after the ‘effective’ transition. For reoccurring patterns (yellow line), IC is expected to show an earlier drop, and these are hence associated with faster RTs (‘RT advantage’).

Reaction times in Barascud et al (2016) were consistent with those obtained from an ideal-observer model based on prediction by partial matching (PPM), as implemented in the IDyOM model (Pearce, 2005, 2018). IDyOM, shown to be an effective model of human auditory sequence learning on multiple time scales (e.g.: Agres, Abdallah, & Pearce, 2018; Harrison & Pearce, 2018; Pearce, 2018; Pearce & Wiggins, 2006), proposes that listeners acquire an internal representation of the sound input through partitioning the stimulus sequence into n-grams of increasing order that are thereon stored in memory; this context is then used to evaluate the expectedness of ensuing sounds by deriving a measure of surprisal (information content or negative log probability). RAN and REG sequences differ in unexpectedness (high for RAN, low for REG). The transition from a random to a regular pattern (RANREG stimulus) can therefore be detected as a salient drop in information content in the model output (Fig. 1), which reflects increasing compatibility between the incoming sounds and the stored context. The pattern of behavioural reaction times as well as brain response latencies recorded from naïve, passively listening participants (Barascud, Pearce, Griffiths, Friston, & Chait, 2016; Southwell et al., 2017; Southwell & Chait, 2018) suggest that listeners indeed identify the emergence of regularity by detecting the associated drop in information content and that such tracking of instantaneous expectedness constitutes an automatic, inherent aspect of auditory sequence processing.

The present study uses a combination of behavioural manipulation and modelling to examine the durations over which these memory representations are maintained by introducing rare pattern reoccurrences. One might expect that detection of regularities benefits not only from immediate sequence context, but also from traces accumulated over a longer period. Participants listened to RAN and RANREG sequences (as shown in Fig. 1), and were instructed to press a keyboard button as soon as possible when a transition to REG was detected. All stimuli were created from a pool of 20 frequencies that occurred with equal probability and roughly equal temporal density in all conditions. The only factor distinguishing different sequences was the specific sequential arrangement of tone-pips over time. New sequences were generated on each trial, but a few different regular patterns reoccurred very sparsely (every ∼2.7 minutes) across trials and blocks (RANREGr). Critically, the RAN portion of those trials remained novel. For each participant, different regularities were designated as reoccurring patterns. Stimuli were presented in blocks of approximately 8 minutes each. Within each block, each RANREGr type trials reoccurred 3 times (about 5% of the trials within a block) and was flanked by many novel patterns (RAN and RANREG).

We hypothesized that, if the stored representation of a pattern strengthens through repetition, the information content associated with a transition to a familiar regularity will dip earlier than that associated with a novel regular pattern (Fig. 1, yellow line in the cartoon model), reaching the putative detection threshold more quickly. Behaviourally this process should be revealed as faster reaction times to recurring patterns (‘RT advantage’ in Fig. 1). The size of this effect may provide a window into the latent variables associated with the retention of sensory information in memory.

Several properties render this paradigm attractive. First, all sequences consist of the same 20 frequency ‘building blocks’. This simplifies parametrization and modelling of the task, while retaining sufficient pattern complexity (there are more than a trillion permutations of 20 frequencies). Second, these 20 frequencies are isochronous and occur with equal probability: stimuli are thus matched in terms of long-term spectrum, average statistics and time patterning. The only difference between RAN and REG patterns and, importantly, between REG and REGr patterns is the specific arrangement of these tone-pips over time. To distinguish a familiar regularity from a novel one, listeners must retain in memory the specific permutation of the tone-pips (we confirm this explicitly in Experiment 1B). Third, the task does not require listeners to *explicitly* memorize sounds (the emergence of the REG pattern readily pops out perceptually; see stimulus examples in sup materials). It thus taps the process by which we automatically glean acoustic information from an ongoing stream so as to detect (a posteriori) regularities.

Across the experiments presented here, we ask whether human listeners form long term memories of sparsely reoccurring regular patterns (yes), whether this memory is robust to interference (yes) and whether it can be formed through passive exposure (partially). Through a combination of behavioural manipulation and modelling we also demonstrate the interplay between short (a few seconds) and long (over many minutes) integration in the process of long-term memory formation. Overall the results reveal a robust, implicit and long-lasting memory for sequential structure. The findings highlight the attunement of the auditory system to exceedingly sparse reoccurrence of sequential acoustic regularities, even when presented within highly similar patterns. This sensitivity may underlie the extensively documented human ability to rapidly learn repeating structure in sensory sequences (Conway & Pisoni, 2008).

## Results

Participants listened to RAN, RANREG, RANREGr, CONT and STEP sequences as illustrated in Fig. 1 and were instructed to monitor for transitions. The reaction time (RT) to STEP was subtracted from the RT to RANREG to estimate a lower bound measure of the time required to detect the emergence of regularity. RT values reported below are all baselined RTs (the raw RTs from which the RT to the STEP condition was subtracted).

Compared with RTs to the emergence of novel regularities (RANREG), we expected progressively faster RTs as regularities reoccur across the experiment (RANREGr), indicating that their representations have become retrievable from memory. We assess overall memory formation of REGr based on RTs averaged over all three reoccurrences within each block. However, we focus on RTs in each intra-block presentation to assess persistence of memory effects across experimental manipulations.

### Experiment 1A: Implicit long-lasting memory for 3 reoccurring patterns

Fig. 2 (A-D) plots the mean and individual results of the regularity detection task performed in three sessions: 5 blocks on day 1, one block after 24 hours (‘24h’) and one block after 7 weeks, (‘7w’). Participants were highly accurate in detecting regularities (Fig. 2A): d’ plateaued at near ceiling performance after the first block. No difference was observed between hit rates for RANREG and RANREGr [no main effect of condition: F(1,19) = .39, p = .539, η_p_^2^ = .02; no main effect of block: F(5, 90) = .46, p = .804, η_p_^2^ = .02; no interaction between condition and block: F(5, 90) = 1.10, p = .367, η_p_^2^ = .06].

**Fig. 2.**
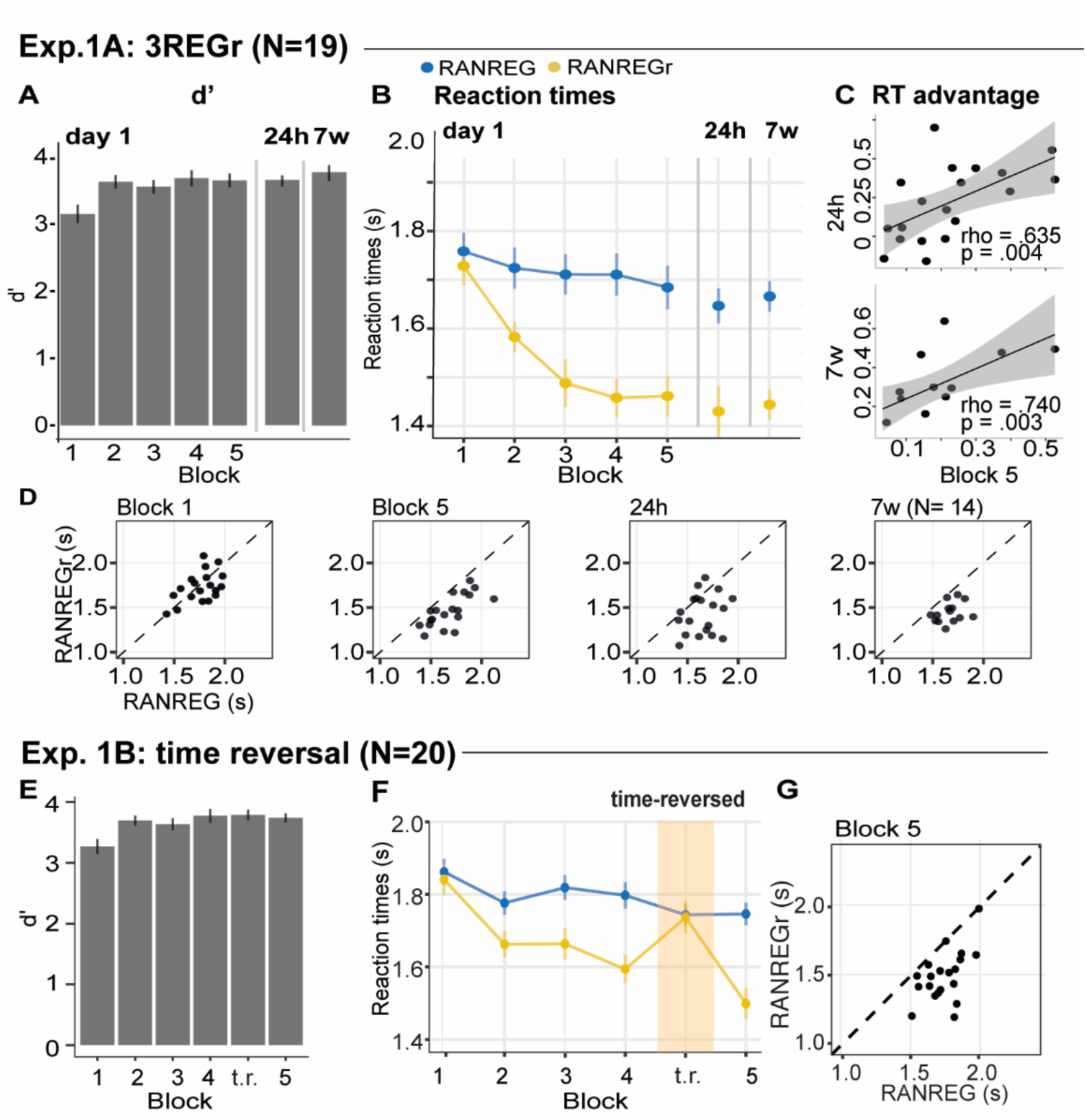
Experiment 1A, B: Implicit long-lasting memory for 3 reoccurring patterns, and specificity to sequential structure. **(A-D) Exp. 1A (3 reoccurring targets). (A)** Sensitivity to emergence of regularity (d’) across blocks during the first session, as well as after 24 hours and after 7 weeks. Error bars indicate 1 s.e.m. **(B)** RT to the transition from random to regular pattern in RANREG and RANREGr conditions, across blocks. Error bars indicate 1 s.e.m. **(C)** Correlations between RT advantage at the end of the first day – block 5 – and after 24 hours (upper plot) and after 7 weeks (lower plot). Each data point represents an individual. Note N=14 in the 7W data due to attrition **(D)** The relationship between RTs for the RANREG and RANREGr conditions. Each data point represents an individual participant. Dots below the diagonal reveal faster detection of RANREGr compared with RANREG. **(E-G) Exp. 1B (time reversal): (E)** Sensitivity to emergence of regularity (d’) across blocks. **(F)** RT to the transition from random to regular pattern in RANREG and RANREGr conditions, across blocks. The block containing time-reversed REGr is shaded in yellow. **(G)** The relationship between RTs to the RANREG and RANREGr conditions in block 5.

Despite the ceiling effects associated with pattern detection (mean hit rate = 97.3 %), faster RTs in the RANREGr than in the RANREG condition (‘RT advantage’) were observed in all participants by the end of the first session (block 5; Fig 2D), indicating a clear implicit memory for the reoccurring patterns. A repeated measures ANOVA on RTs with condition (RANREG and RANREGr) and block as factors yielded a main effect of condition [F(1,18) = 34.09, p < .001, η_p_^2^ = .65], main effect of block [F(5,90) = 9.24, p < .001, η_p_^2^ = .3] and an interaction between condition and block [F(5,90) = 6.88, p < .001, η_p_^2^ = .28]. Specifically, in the first block of the first session, performance did not differ between RANREG and RANREGr [t(18) = .794, p = 1]. By the end of the second block (after 6 REGr reoccurrences), a significant difference (∼ 140 ms; 2.8 tones) between RTs was observed [REG –REGr: t(18) = 3.964, p = .006]. This difference grew over the following blocks (all ps < .001), plateauing after block 3 (233±.17 ms; 4.7 tones). The RT advantage on the third block did not differ from the fourth [t(18) = −0.907, p = 1] nor from the fifth block [t(18) = −.0003, p = 1]). In **Experiment S1** (Fig. S4), we demonstrate that similar effects are obtained when doubling the number of REGr patterns to be memorised (6 different patterns per participant). In **Experiments S2A and S2B** (Fig. S5), we further demonstrate that the memory trace is not abolished by introducing ‘interrupting blocks’ (in which the RANREGr condition was not presented) between ‘standard blocks’ (in which RANREGr patterns reoccurred every ∼2.7 minutes).

Critically, implicit memory for reoccurring regularities persisted after 24 hours and after 7 weeks: the RT difference between novel and reoccurring sequences remained constant between the last block of day 1 (block 5) and after 24 hours [t(18) = .139, p =.891], as well as between 24 hours and 7 weeks later [t(13) = −.668, p =.515]. An inspection of intra-block reoccurrences (Fig. S1A) revealed that the RT advantage for REGr was similar between the third (last) intra-block presentation of day 1 and the first intra-block presentation after 24 hours [t(18) = .123, p =.903]; similarly, in the session conducted after 7 weeks, the RT advantage measured after the first intra-block presentation was similar to the third (last) presentation in the session conducted after 24 hours [t(13) = .958, p =.356; (Fig. S1A)]. This suggests that the effect observed after 24 hours and 7 weeks reflects the presence of a lasting memory trace of reoccurring regularities rather than rapid within-block re-learning.

Further, we examined the correlation of individual participants’ RT advantage across the three sessions (Fig. 2C). A robust correlation was found between the end of the first day (block 5) and the measurement taken after 24 hours (spearman’s rho= .635, p = .004) – participants who exhibited a large RT advantage at the end of the first day were also those showing a larger advantage 24 hour later. A similar correlation was found with performance after 7 weeks (spearman’s rho= .740, p = .003). This confirms strong reliability of individual effects.

### The memory effects are not driven by explicit recognition of reoccurring patterns

Explicit memory for reoccurring regularities was examined at the end of each session by means of a familiarity task. Only regular sequences were presented: REGr (one presentation of each pattern) were intermixed with previously unheard REG patterns. Participants were instructed to indicate which patterns sounded ‘familiar’. Classification was evaluated using the MCC score (see methods) which ranges between 1 (perfect classification) to −1 (total misclassification). The mean MCC on day 1 was 0.231 (see Fig. S2A) and this measure did not correlate with the RT advantage observed in block 5 (spearman’s Rho=0.307; p=0.201; a similar result was also obtained when pooling across participants from Exp. 1A and Exp. S1 (which used 6 REGr patterns) (Spearman’s Rho=0.114; p=0.493; N=38). Therefore, implicit memory for reoccurring patterns, observed in nearly all participants, is not linked to explicit awareness of reoccurrence. In line with this, though a weak correlation between RT advantage and MCC measured after 24 hours (Spearman’s Rho= .459, p = .048, N=19), there was none after 7 weeks (Spearman’s Rho= −.024, p = .934, N=14).

### Experiment 1B: Implicit memory is specific to sequential structure

To confirm that the RT advantage effects are driven by memory of sequential structure, in this experiment (Fig. 2E-G) we tested whether implicit memory for reoccurring patterns is tolerant to time reversal of the originally learned patterns. Participants performed the regularity detection task as in Exp. 1A over 6 experimental blocks. The first 4 were identical to those in Exp. 1A. In the fifth block, REGr sequences were replaced by time-reversed versions. In block 6, the original REGr were introduced again. Participants were naïve to the experimental manipulation. It was expected that, if implicit memory is specific to the sequential structure of regularity, the RT advantage should disappear in the time-reversed block (see also Kang et al, 2017).

Blocks 1-4 revealed the same effects as in Exp. 1A (Fig. 2F)[main effect of condition: F(1,19) = 71.96, p < .001, η_p_^2^ = .79; main effect of block: F(3, 5) = 9.90, p < .001, η_p_^2^ = .34; interaction condition by block: F(3, 57) = 5.67, p < .001, η_p_^2^ = .23]. Specifically, in the first block RTs in the RANREGr condition were similar to those in RANREG [t(19) = 0.725, p = 1], but became progressively faster (114 ms; 2.27 tones) in the second block [t(19) = 3.56, p = .01], and across the remaining blocks (all ps < .001) (203 ms; 4.1 tones in the 4^th^ block).

Importantly, this RT advantage was abolished in the time-reversed block, but restored in the subsequent block containing the originally learned REGr: a repeated measures ANOVA with condition (RANREG and RANREGr) and the last two blocks as factors yielded a main effect of condition (F(1,19) = 25.57, p < .001, η_p_^2^ = .57), a main effect of block (F(1, 19) = 18.09, p < .001, η_p_^2^ = .49), and an interaction condition by block (F(1, 19) = 40.03, p < .001, η_p_^2^ = .68), demonstrating the significantly greater RT advantage (RANREG novel – RANREGr) in the last than in the time-reversed block [t(19) = 6.33, p < .001]. The RT advantage for REGr in the third intra-block presentation of block 4 (Fig. S1B) was greater than in the first intra-block presentation of the time-reversed block [t(19) = −2.261, p =.035], but similar to the first intra-block presentation of the last block reintroducing the original REGr [t(19) = .788, p =.440].

These results constrain the nature of the observed memory effect to sequential information.

### Experiment 2: Limited formation of memory traces of non-adjacent patterns

In this experiment (Fig. 3) we tested whether *adjacent* repetition of patterns (as is inherently the case for REG sequences) is required for implicit memory to be formed.

**Fig. 3.**
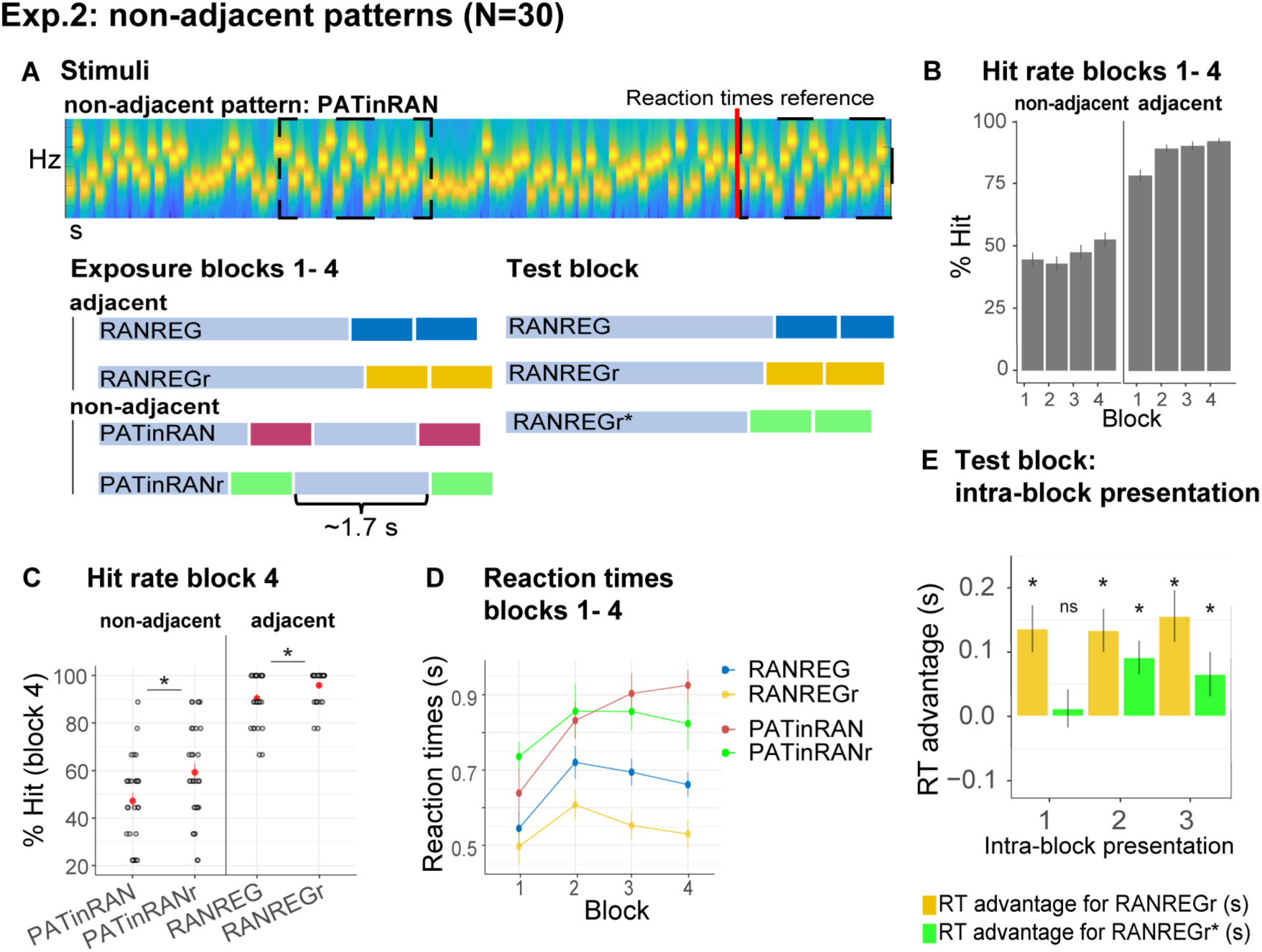
Experiment 2: Limited formation of memory traces of non-adjacent patterns. **(A)** In blocks 1 to 4, listeners were exposed to RAN, RANREG, RANREGr, PATinRAN and PATinRANr trials. An example spectrogram for a PATinRAN stimulus is provided. The non-adjacent repetitions of the 20-tones pattern (PAT) are indicated by dashed rectangles. In block 5 (‘test’ block’) PATinRANr sequences were replaced by versions where the 2 cycles were set adjacent at the end of the trial (RANREGr*). **(B)** Accuracy (block 1 to 4): hit rates are computed separately for adjacent (RANREG and RANREGr) and non-adjacent (PATinRAN and PATinRANr) trials. **(C)** Hit rates in block 4, separately for novel and reoccurring adjacent and non-adjacent conditions. ‘*’ indicates a significant difference between conditions. **(D)** RT (measured relative to the onset of the second cycle; see red line in A) across blocks 1 to 4 for RANREG, RANREGr, PATinRAN and PATinRANr. Error bars indicate 1 s.e.m. Note that since RT here is computed relative to the onset of the REG repetition, to compare RANREG RT with those reported in figures above, add 1 sec **(E)** Test block: RT advantage for RANREGr (yellow) and RANREGr* (green) in each intra-block presentation. Error bars indicate 1 s.e.m. To determine the presence of a memory trace to REGr* we specifically focus on the first intra-block presentation. ‘*’ indicates a significant RT advantage, ‘ns’ indicates an RT advantage not significantly different from 0.

Over 4 blocks, listeners were exposed to RAN, RANREG and RANREGr trials as in previous experiments. We also introduced a new condition, PATinRAN (Fig. 3A), which consisted of two identical *non-adjacent* 20-tone patterns (PAT) embedded within a random sequence of tone-pips. The second appearance always occurred at the end of the sequence. The first appearance was embedded partway through the sequence at an average distance of 1.7 seconds (range 0.5-2.9 s). To understand whether memories of non-adjacent patterns (PAT) can be formed during listening, 3 different PAT reoccurred 3 times within block (PATinRANr; the random parts of the sequences as well as the separation between the two PAT patterns remained random on each trial).

Both non-adjacent (PATinRAN, PATinRANr) and adjacent (RANREG, RANREGr) trials included two repetitions of each pattern with the only difference being that they were contiguous in the latter and separated by random tones in the former. Participants were instructed to respond if they detected two identical, not necessarily contiguous, 20-tone patterns within a trial; 50% of the trials consisted of fully random patterns. In order to make sure that participants paid equal attention to the (harder) PATinRAN sequences, accuracy was emphasized over response speed.

In the last block (block 5; ‘test block’), we tested whether, following a comparable amount exposure through block 1 to 4, PATinRANr and RANREGr patterns were similarly remembered. To equate difficulty of pattern detection in this block, aside from any learning-related difference, PATinRANr sequences were replaced by versions where the 2 cycles were set adjacent at the end of the trial. We refer to these conditions as RANREGr*. Participants were instructed to respond as quickly as possible. We compared the magnitude of the RT advantage associated with RANREGr* to that associated with RANREGr.

Fig. 3B shows the detection performance during the exposure blocks (1 to 4). Despite having practised the PATinRAN condition, detection performance was overall worse, and substantially more variable in PATinRAN (mean over blocks 1-4: 47.36 ± 16.5 %) relative to RANREG (88.47 ± 11.6 %), and improved less across blocks [main effect of condition: F(1, 29) = 419.01, p < .001,, η_p_^2^ = .94; main effect of block: F(3, 87) = 9.24, p < .001, η_p_^2^ = .24; interaction of condition per block: F(3, 87) = 4.83, p = .004, η_p_^2^ = .14]. Thus, whilst a pattern is highly detectable when contiguously repeated, performance drops substantially when the repetition is not adjacent, presumably due to limits on short-term memory.

Focusing on the 4^th^ block (Fig. 3C): A repeated measures ANOVA with the factors reoccurrence (novel / reoccurring patterns) and adjacency (adjacent / non-adjacent patterns) yielded a significant main effect of adjacency [F(1, 29) = 205.99, p < .001, η_p_^2^ = .88]: as expected, whilst participants were very apt at detecting RANREG patterns, performance on PATinRAN was substantially more variable and lower overall. Interestingly a main effect of reoccurrence [F(1, 29) = 21.74, p < .001, η_p_^2^ = .43], was also observed, with no interaction between the two factors [F(1, 29) = 3.95, p = .056, η_p_^2^ = .12]. Therefore, detection data showed an increase in accuracy for reoccurring patterns in both adjacent and non-adjacent conditions. The emergence of this effect for RANREGr, despite its absence in Exp. 1A, is presumably driven by the below ceiling performance observed here (mean hit rate = 93% relative to 97.5 % in Exp. 1A) – likely a consequence of the extra behavioural strain introduced by the PATinRAN stimuli. Critically, the finding of increased hit rates for PATinRANr (a mean increase of 15%) demonstrates that, through repeated exposure, listeners formed a memory trace for the non-adjacent patterns.

RT results across block 1 to 4 are shown in Fig. 3D. To allow for a comparison across conditions, RTs here are measured relative to the onset of the second regularity cycle (indicated with a red line in Fig. 3A). Since participants were encouraged to prioritise accuracy over speed in these blocks, the RT data in blocks 1-4 were not statistically analysed. However, an RT advantage (reaching 131 ms –2.63 tones in block 4) is clearly visible for RANREGr relative to RANREG stimuli.

Test block: as a critical test for the formation of memory traces, we assessed the presence of a RT advantage in the 1^st^ intra-block presentation of RANREGr and RANREGr* (Fig. 3E). The RT advantage was significantly different from zero in RANREGr [one-sample t-test: t(29) = 3.724, p = .001], but not in the RANREGr* condition [one-sample t-test: t(29) = .419, p = .678]. A paired t-test further confirmed a greater RT advantage in the RANREGr than in the RANREGr* condition [(t(29) = 3.169, p = .003]). This indicates that, as a group, participants did not demonstrate an immediate RT advantage to RANREGr* patterns. As seen in Fig. 3E, a RT advantage in RANREGr* emerged following the second intra-block presentation. This effect may be associated with learning within the test block. A repeated measures ANOVA on RT advantage in the test block with the factors condition (REGr/ REGr*) and intra-block presentation (1st/ 2nd/ 3rd) revealed a main effect of condition [F(1, 29) = 9.09, p = .005, η_p_^2^ = .24] but no main effect of intra-block presentation [F(2, 58) = .67, p = .515, η_p_^2^ = .02], or interaction [F(2, 58) = 1.27, p = .287, η_p_^2^ = .04], consistent with an overall smaller RT advantage to RANREGr*.

As an exploratory analysis, we tested whether higher detection accuracy for non-adjacent patterns (hit rates in block 4 for PATinRANr /PATinRAN) predicted a greater RT advantage when the patterns were set adjacently in the test block (REGr*). We observed a significant moderate correlation between the detection accuracy of PATinRANr in block 4 and the RT advantage in the 1^st^ intra-block presentation of REGr* (spearman’s rho= .429, p = .018) such that those participants who exhibited a higher detection accuracy for PATinRANr in block 4, were also associated with a higher RT advantage for REGr* in the test block. This correlation with RT advantage was specific to PATinRANr, in that it did not extend to PATinRAN (spearman’s rho= .017 p = 0.927) and held when the effect of detection accuracy for PATinRAN was accounted for (spearman’s rho = .465, p = .011). The specificity to PATinRANr suggests that the link is not simply related to some property of short-term memory (in which case we would have expected a correlation with PATinRAN as well) but is specific to the memory advantage for PATinRANr stimuli which developed over the first 4 blocks.

Overall, these results suggest the presence of limited memory traces for reoccurring, non-adjacent patterns (PATinRANr). However, it is clear that the formation of robust implicit memory traces for sound sequences depends on short-term memory (and hence benefits from immediate repetition of patterns) such that introducing a gap of even 2 seconds results in substantially weakened storage in memory.

### Modelling

We constructed a computational model, based on ‘prediction by partial matching’ (PPM; see Methods) to provide a formal simulation of the psychological mechanisms underlying the behavioural effects in Experiments 1A (Fig 2), 2 (Fig. 5) and S2A (Fig. S5D). These experiments reflect critical manipulations of the effect of long- and short-term memory. The model yielded a single set of values which best account for listeners’ performance across those experiments.

To model the present behavioural data, the computational model instantiates the following cognitive hypotheses:

*1)* Listeners learn sequence statistics throughout the experiment. This statistical learning takes the form of *n*-gram memorisation. The more a listener is exposed to a given *n*-gram, the stronger salience this *n*-gram has in memory. Here, we allow *n* to range between 1 and 5, corresponding to sub-sequences of 1 and 5 tones respectively.
*2)* The listener uses these *n*-gram statistics to quantify the predictability of incoming tones based on the preceding portion of the sequence and other information stored in memory as a generative probabilistic model (represented by PPM, see Methods).
*3)* Listeners monitor the information content (IC; negative log probability) of tones throughout the tone sequence, where high IC corresponds to low probability and low IC corresponds to high probability. Sudden changes in IC are indicative of potential changes in the environment. In the present case, a sudden drop in IC reflects the onset of repetitive structure in the stimulus corresponding to a transition from RAN to REG. Once the model is sufficiently confident that a reliable drop has occurred, it registers a ‘change detected’ response analogous to the participant’s button press.
*4)* The memory salience of a given n-gram observation decays over time, with this decay profile reflecting the dynamics of human auditory memory. In particular, we adopt the memory-weighting scheme recently presented in Harrison, Bianco, Chait, & Pearce (2020), and implement the following decay profile for the memory salience of an n-gram observation: a) an initial short and high-fidelity steady-state phase, representing an echoic memory buffer; b) a fast exponential-decay phase, representing short-term memory; c) a slow exponential-decay phase, representing longer-term memory (see Fig. 4A, Table 1 for more details). The model also adds noise to the memory retrieval stage, simulating inaccuracies in human memory retrieval.

**Fig. 4.**
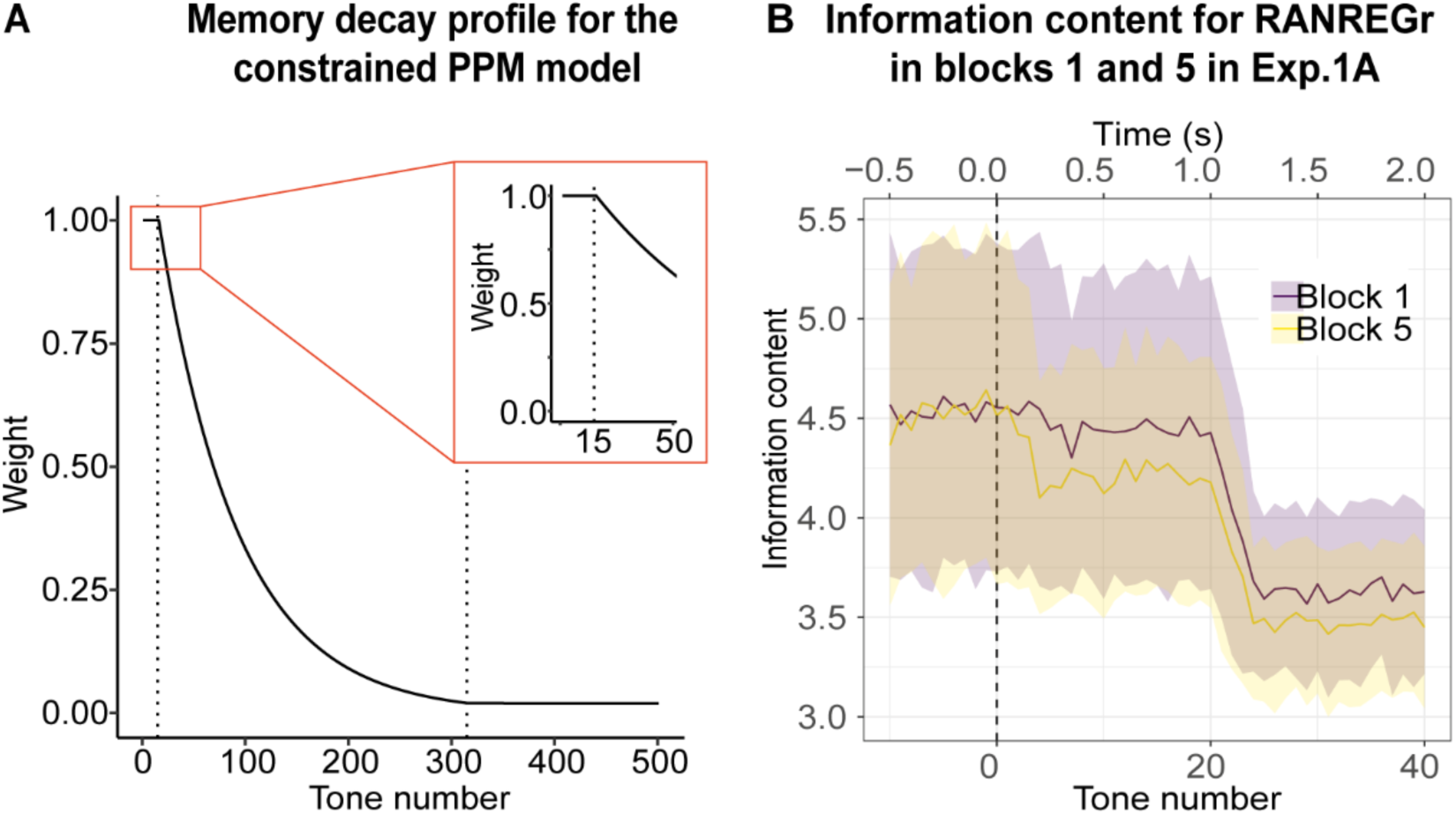
Memory constrained PPM model. **(A)** Memory decay profile for the constrained PPM model. The curve describes the weight of a given n-gram observation in memory as a function of the number of consequent tones that have been presented, assuming a constant presentation rate of 20 Hz. The two dotted lines indicate transitions between the different phases of memory decay: the first, between the memory buffer and short-term memory, and the second, between short-term memory and long-term memory. The inset shows the transition from the memory buffer (of 15 tones capacity) to the fast exponential-decay phase. See Table 1 for model parameters. **(B)** Information content as a function of tone number for RANREGr trials in blocks 1 and 5 of Exp. 1A. Mean Information content is computed from the memory-decay PPM model, expressed in bits, and averaged over all trials. The shaded ribbons correspond to 1 STDEV. Trials are aligned such that a tone number of 0 corresponds to the first REG tone after the transition. The transition between RAN and REG phases becomes clearest after about 24 tones; however, the model detects the transition faster in block 5 than in block 1, because it partially recognises the REGr cycle from its previous occurrences, yielding a lower information content that is more clearly distinguishable from the RAN baseline and therefore requires less evidence accumulation time (= faster detection). However, it is obvious from the large error bars that the effects are subtle.

**Fig. 5.**
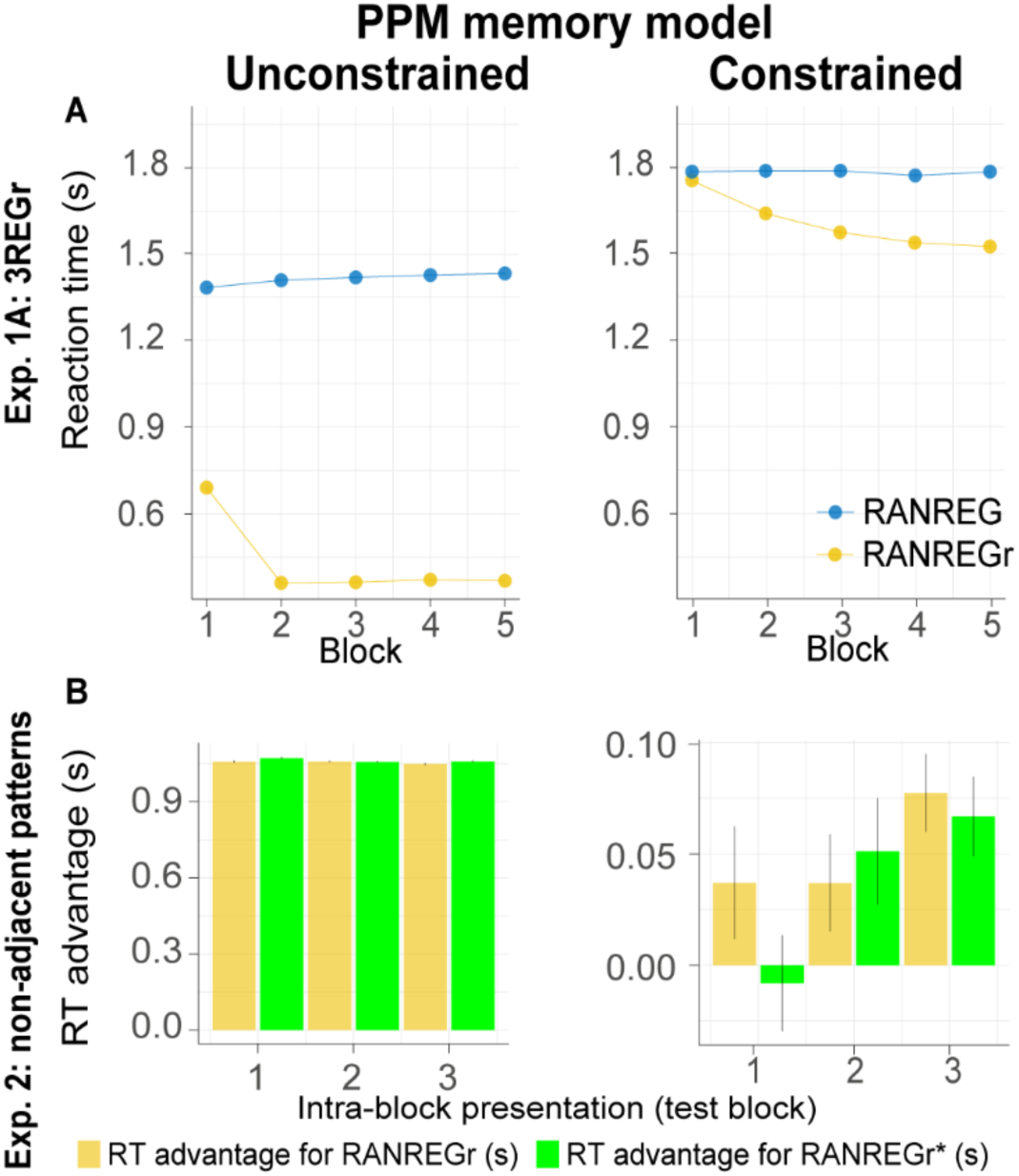
Model simulations for Experiments 1A and 2 for the unconstrained (left) vs. constrained (right) PPM model. Overall, we demonstrate a qualitative similarity between the formal simulation of constrained memory and human responses. **(A) Exp. 1A:** the estimated RTs to the transition from random to regular patterns in RANREG and RANREGr conditions across 5 consecutive blocks. The unconstrained model detects the novel RANREG transitions after about 1.4 seconds (18 tones). RANREGr trials in contrast, are associated with substantially quicker detection time which emerges rapidly. Human performance is better accounted for by the constrained model, whereby the detection advantage for RANREGr trials grows over successive presentations of the REGr patterns. Error bars indicate 1 s.e.m. **(B) Exp. 2:** RT advantage in RANREGr and RANREGr* conditions relative to novel RANREG for each intra-block presentation within the test block. Data are presented in the same way as those in Fig 3E. The unconstrained model reveals an equal RT advantage in both conditions. Conversely, similar to human performance, the constrained memory model does not learn the reoccurring non-adjacent patterns across blocks 1 to 4, as shown by the null RT advantage in the first intra-block presentation in the RANREGr* condition. RTs of 1^st^, 2^nd^ or 3^rd^ intra-block presentations were averaged across the different RANREGr/RANREGr*, and RTs to RANREG novel were averaged across trials which occurred at the beginning (first third), middle or end of each block. Error bars indicate 1 s.e.m.

**Table 1.** Parameters for the memory-decay PPM model as manually optimized for Experiments 1A, 2, and S2A. *The combination of STM weight, STM duration and LTM weight yields a STM half-life of 3.06 seconds.

| Parameter | Value |
| --- | --- |
| Buffer capacity | 15 items |
| Buffer weight | 1 |
| Short-term memory weight* | 1 |
| Short-term memory duration* | 15 seconds |
| Long-term memory weight* | 0.02 |
| Long-term memory half life | 500 seconds |
| Long-term memory asymptote | 0 |
| Noise | 1.3 |
| Order bound | 4 |

As a benchmark, we also report the results for an equivalent unconstrained model (i.e., with perfect memory). Although the parameters shown in Table 1 were not formally optimised to fit the empirical data quantitatively, the memory constrained model shows close qualitative correspondence to the pattern of RTs observed in Experiments 1 and 2 in a way that the unconstrained model clearly does not (see Fig. 5). Note that the change point detection algorithm was configured with a strict threshold in order to achieve an appropriate Type I error rate (see Methods).

In **Experiment 1A**, participants experienced progressive facilitation for the RANREGr trials over the course of the experiment’s five blocks, reflecting accumulated long-term memory traces for tone patterns shared across trials. Fig. 5A (left) simulates Exp. 1A using an unconstrained PPM model. For RANREG trials, the REG patterns are novel for each trial, and hence the repeating cycles can only be detected once one complete cycle has elapsed and the sequence embarks on the second cycle. For these trials, the unconstrained PPM model detects transitions after about 1.4 seconds, corresponding to one complete cycle plus eight tones. In contrast, for RANREGr trials after the first stimulus block, the regular patterns are already familiar from previous trials. The unconstrained PPM model remembers these previous trials perfectly, and hence experiences a sharp drop in IC soon after the beginning of the repeated cycle, corresponding to a reaction time of about 0.4 seconds or 8 tones. In contrast, human participants experienced a much slower and less extreme facilitation.

A better account of Exp. 1A can be achieved by introducing memory decay into the PPM model. Fig. 5A (right) shows the reaction-time profiles simulated by the constrained memory model. Like the human participants, the model experiences a moderate facilitation effect that grows over successive presentations of identical regular patterns. Fig. 4B illustrates this effect in more detail, plotting average information content profiles for RANREGr trials in Block 5 as compared to RANREGr trials in block 1.

It is important to note that the steady long-term decay, which is a key feature of the memory constrained model predicts that the performance facilitation should disappear after 24 hours, and certainly after 7 weeks. After such time periods, the memory traces for the reoccurring patterns should decay to zero, and the corresponding facilitation effect should disappear. Remarkably, the participants exhibited unaltered performance facilitation. This suggests that the memory traces of these reoccurring patterns are somehow ‘fixed’ at a certain point during testing (perhaps after about 3 blocks; see Exp. 3 below). One way of simulating this effect would be to change the asymptote of the exponential memory decay, such that the memory trace asymptotically approaches a small but non-zero value as time tends to infinity. However, we found that incorporating such an asymptote caused the performance facilitation for RANREGr trials to increase constantly from block to block, in contrast with the plateau in performance facilitation shown in the behavioural data after about three blocks. It seems likely, therefore, that there remains a non-trivial ‘fixing’ effect that is not accounted for by the current model.

**Experiment 2** investigated the effect of pattern adjacency on pattern detection and memory formation. We trained unconstrained and constrained models on blocks 1-4, and report their performance for block 5, the ‘test’ block where PATinRANr sequences were replaced by versions where the 2 non-adjacent cycles were set adjacent at the end of the trial (RANREGr*). The RANREG and RANREGr conditions were preserved from blocks 1-4. Block 5 therefore provided a direct test of whether pattern learning was influenced by the adjacency of the target patterns in blocks 1-4. The unconstrained PPM model is unaffected by adjacency (Fig. 5B left): the immediate advantage shown at the 1^st^ intra-block presentation for both RANREGr* and RANREGr conditions shows that the unconstrained-memory model learns just as well from non-adjacent cycles as from adjacent cycles. However, the behavioural data indicate that the human participants were indeed impaired by non-adjacency (Fig. 3F). The memory-decay PPM model (Fig. 5B right) fully reproduces this pattern of results.

### Experiment 3: Memories of a set of reoccurring regularities are not overwritten by subsequent memorization of another set

Does memorization of a new set of REGr interfere with the representation of a previously memorized set? Participants performed the same transition detection task as in Exp. 1A. They were exposed to a set of 3 reoccurring patterns (REGr1) in the first 3 blocks, followed by 3 blocks in which another set of patterns (REGr2) reoccurred. Blocks 7 and 8 then re-tested memory for the reoccurring regularities of set 1 and set 2 respectively. After 24 hours, memory for the two sets of regularities was tested again.

Clear implicit memory for the first set of targets (REGr1), as indicated by an RT advantage, was observed after the 3^rd^ block (Fig. 6B) [main effect of condition: F(1,28) = 41.01, p <.001, η_p_^2^ =.59; main effect of block: F(3,84) = 15.69, p < .001, η_p_^2^ =.36; condition by block interaction: F(3, 84) = 6.83, p < .001, η_p_^2^ =.20]. As expected, after 3 blocks of exposure the RT advantage in the RANREGr1 condition (163 ms – 3.3 tones) was similar to that observed in Exp. 1A above. Critically, this RT advantage for RANREGr1 was not perturbed after the presentation of the second set of regularities (REGr2) [RT advantage in block 3 vs. block 7: t(28) = .877, p =.387]. It also lasted after 24 hours [RT advantage in block 7 vs. after 24 hours: t(28) = −.553, p = .584], and was similar to the 24h RT advantage observed in Exp. 1A [no main effect of experiment: F(1,50) = .33, p =.567, η_p_^2^= .01]. These results indicate that, once formed, memory traces are neither overwritten nor weakened by ‘interfering’ new sets of reoccurring patterns.

**Fig. 6.**
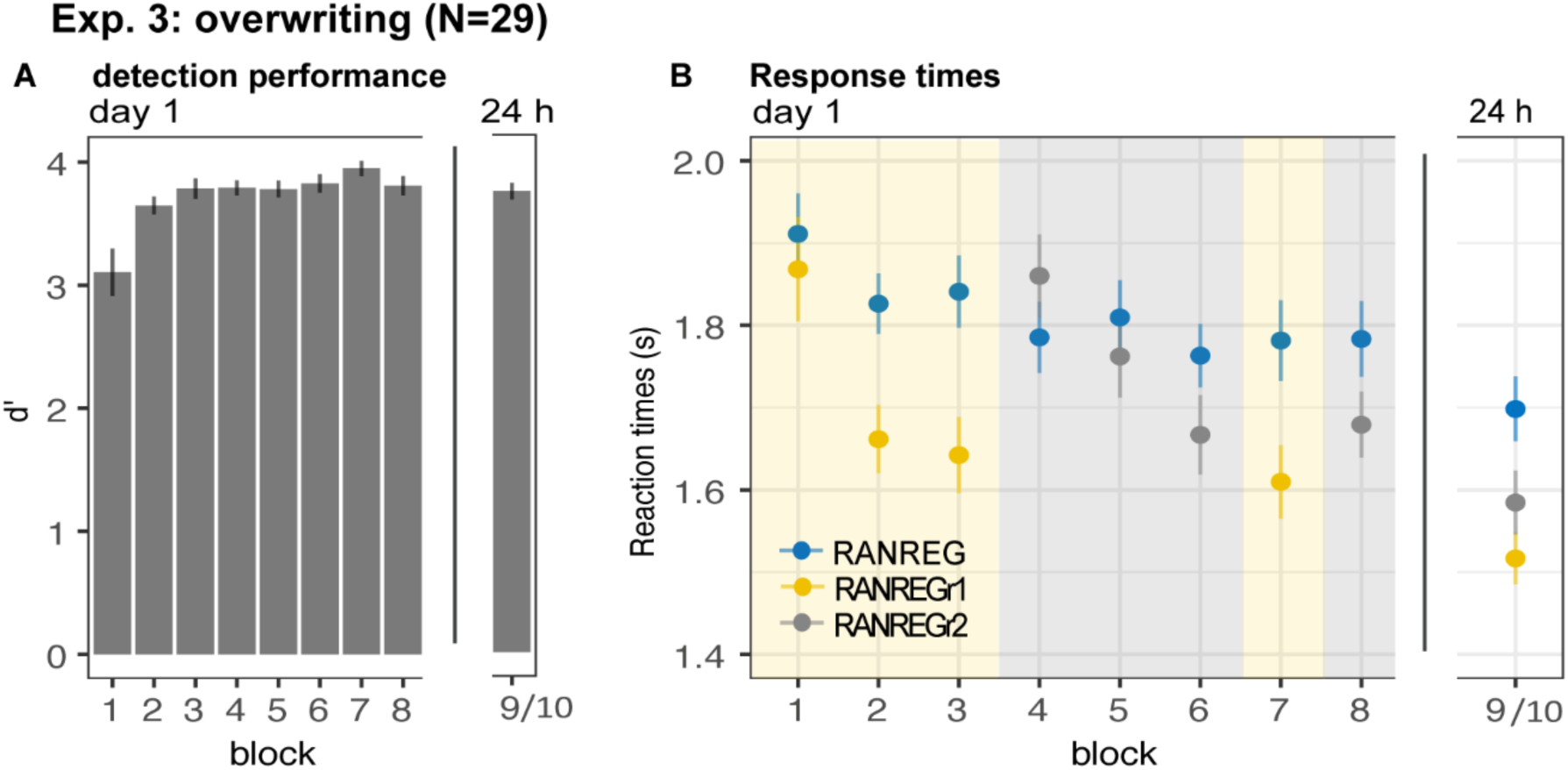
Experiment 3: memories of a set of reoccurring regularities are not overwritten by subsequent memorization of another set. Participants were exposed to a set of 3 reoccurring patterns in the first 3 blocks (REGr1, yellow shading), followed by 3 blocks in which another set of patterns was reoccurring (REGr2, grey shading). The final blocks (7 and 8) tested memory for set 1 and 2, respectively. After 24 hours, memory for the two sets was tested again. **(A)** d’ across all blocks on day 1 and after 24 hours. Error bars indicate 1 s.e.m. **(B)** RT to the transition from random to regular pattern across blocks for RANREG, RANREGr1 and RANREGr2 on day 1 and after 24 hours. Error bars indicate 1 s.e.m.

In blocks 4-6 presenting the second set of reoccurring regularities (REGr2) also showed an RT advantage, as demonstrated by the emerging separation between the RT to novel and reoccurring regularities. A repeated measures ANOVA on the RT advantage with ‘experimental stage’ (blocks 1-3, blocks 4-6) and block number (1^st^, 2^nd^ or 3^rd^) showed a main effect of block number [F(2,56) = 20.13, p < .001, n_p_2 = .42; consistent with a growing RT advantage across blocks], and stage [F(1,28) = 15.70, p < .001, n_p_2 = .36] with no interactions. The main effect of stage suggests an overall larger RT advantage for the first set (REGr1). The noisier overall RT pattern observed in blocks 4-6 may be indicative of an order/fatigue effect. Importantly, at the end of day 1 the RT advantage for the two sets of reoccurring regularities did not differ (block 7 vs. block 8: t(28) = 1.721, p =.096]. The RT advantage for the second set was maintained when tested after 24 hours (RT advantage of last block of day 1 vs. after 24 hours: t(28) = −.277, p = .784), and did not differ from that of the first set [RT advantage after 24 hours for RANREGr1 vs. RANREGr2 t(28) = 1.848, p = 0.075].

### Experiment 4: Implicit memory is robust to pattern phase shifts

In all the previous experiments reoccurring regularities were always presented at the same phase of the REG cycle. Here we asked whether the resulting memory trace was anchored to this fixed boundary – i.e., whether listeners remembered the pattern as a specific ‘chunk’ (Dehaene, Meyniel, Wacongne, Wang, & Pallier, 2015; Thiessen, 2017). If so, the RT advantage should reduce when REGr are phase shifted.

Listeners were presented with 6 reoccurring regularities (REGr) over 3 blocks. In block 4, identical REGr were presented but each presentation was associated with a shifted onset relative to the originally presented pattern (see Fig. 7A, and Methods).

**Fig. 7.**
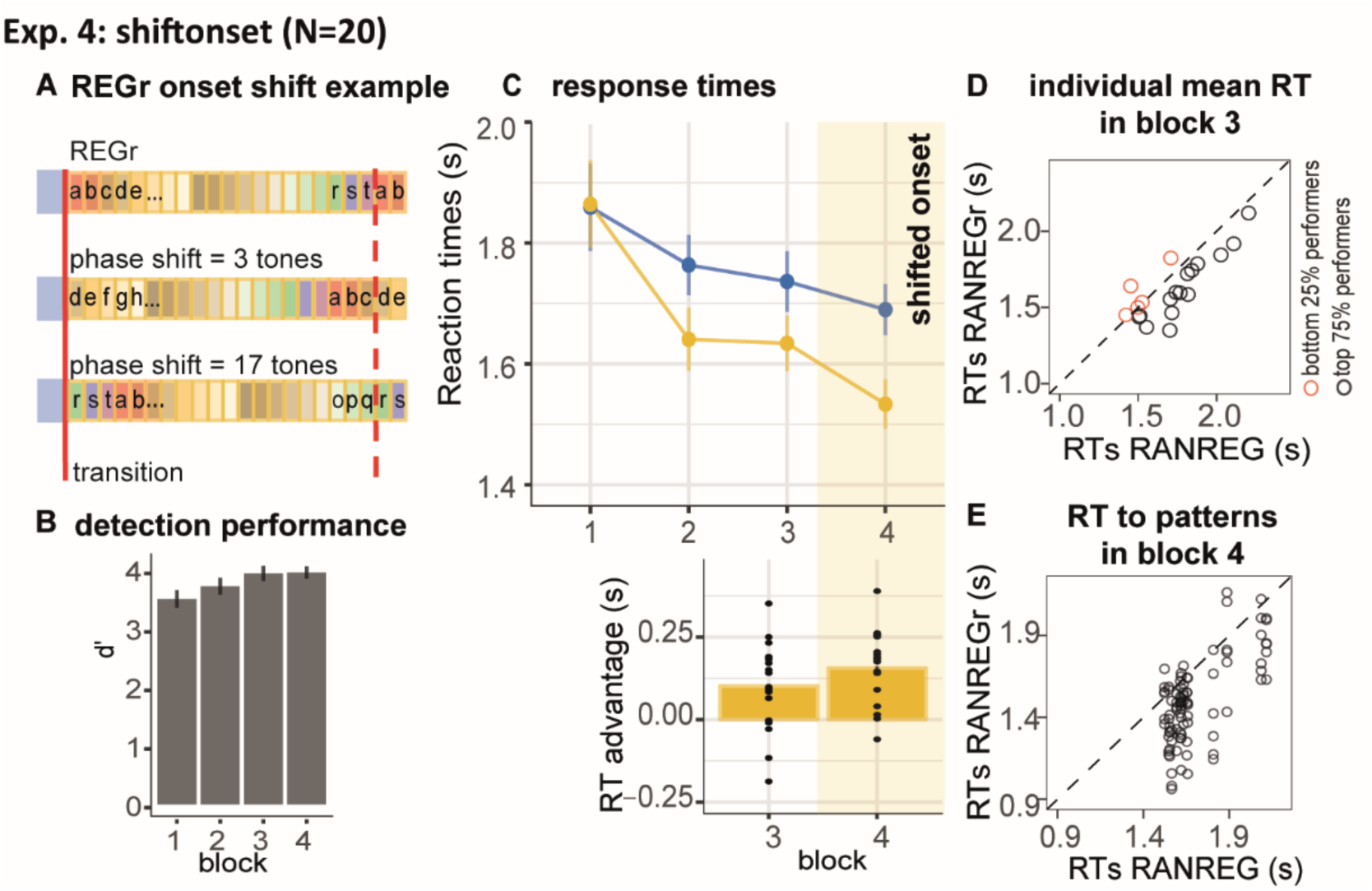
Experiment 4: Implicit memory is robust to pattern phase shifts. **(A)** In this experiment, 6 different reoccurring regularities (REGr) per participant were presented. In block 4 (yellow shading in C) these patterns were replaced by versions with shifted onset of the originally learned REGr. Two examples of phase shifted REGr and their original REGr version are depicted. The solid red line indicates the transition between RAN and REG (the onset of the regular pattern); the dashed red line denotes one cycle (20 tones) **(B)** d’ across all blocks. Error bars indicate 1 s.e.m. **(C)** RT to the transition from RAN to REG pattern across blocks for RANREG and RANREGr. The bottom plot represents the RT advantage observed in blocks 3 and 4. Error bars indicate 1 s.e.m. **(D)** The relationship between RTs to RANREG and RANREGr conditions in block 3. Each circle represents an individual participant. Participants in the bottom quartile (those who exhibited the smallest RT advantage) are shown in red. **(E)** Focusing on those participants who showed learning effects in block 3 (top 75% performers; black circles in D). Plotted is the relationship between RTs to the RANREG and RANREGr in block 4. Each circle represents a unique REGr pattern (6 per participant), plotted against the mean RT to RANREG for that participant., 84.4% of the shifted-onset REGr exhibited a memory effect.

Fig. 7C shows the progressive emergence of the RT advantage associated with the memorization of the reoccurring patterns [main effect of condition: F(1,19) = 21.12, p < .001, η_p_^2^ = .53; main effect of block: F(3, 57) = 18.52, p < .001, η_p_^2^ = .49; condition by block interaction: F(3, 57) = 10.64, p < .001, η_p_^2^ = .36]. Specifically, whilst in the first block performance did not differ between RANREG and RANREGr [t(19) = −.876, p = 1], a faster RT to the RANREGr condition developed across ensuing blocks. This effect continued into block 4, where phase-shifting was introduced (Fig. 7C bottom plot). The RT advantage for phase-shifted RANREGr (167 ms – 3.35 tones) in block 4 was greater than the RT advantage in block 3 (100 ms; 2 tones) [block 3 vs. block 4: t(19) = −13.111, p < .001], demonstrating a strengthening (rather than disappearing) memory effect. The immediate robustness to phase shifting was confirmed by comparing the RT advantage in the first intra-block presentation in block 4, to that in the third (last) intra-block presentation in block 3 (Fig S1C). No significant difference was observed [t(19) = 1.069, p =.298], supporting the conclusion that the RT advantage persisted despite phase shifting.

Further tests confirmed that the RT advantage for REGr in block 4 was similar across small and large phase shifts: a repeated measures ANOVA with factor phase shift (small / large, namely 1-5 and 16-19 vs. 6-15 tones from the original onset) yielded no significant effect of phase shift on the RT advantage [F(1, 19) = .74, p = .400].

These results suggest that sequences are not represented as a fixed chunk of sequential items which is retrieved as a single unit, but more likely as a collection of sequential predictions that are flexibly retrieved from memory according to the available sensory information.

As a further probe into the nature of the representation of the pattern in memory, in **Experiment S3** we investigated listeners’ tolerance to small frequency transpositions. We reveal a transfer of the RT advantage to the transposed pattern, suggesting that the formed representation is not of an exact echoic nature. It is possible that tolerance to frequency transposition reflects a ‘fuzzy’ spectral representation, though we note that the spacing in the present pool – 12% – is generally larger than the just noticeable difference (JND) for frequency typically exhibited by non-musically trained listeners (Tervaniemi, Just, Koelsch, Widmann, & Schröger, 2005). Alternatively, the tolerance to transposition may suggest that instead of the specific frequency pattern, the auditory system maintains a representation of the contour, or inter-tone interval within the pattern.

### Experiment 5: Implicit memory can form during passive exposure, but does not immediately transfer to behaviour

We asked whether memories for reoccurring patterns are formed when sequences are not behaviourally relevant. Naïve participants were exposed to three blocks of the same kind as in Exp. 1A, but instructed to detect the STEP changes only and ignore the other sounds. In the fourth block (‘test’ block) they were instructed to also detect the RANREG transitions.

Sensitivity to transitions in the test block (Fig. 8A) was high overall (mean d’ = 2.77 ± .73), but lower than in the first block of Exp. 1A [independent sample t(35) = −2.140, P =.039]. This is likely because, in order to keep them naïve, participants did not receive training on RANREG detection.

**Fig. 8.**
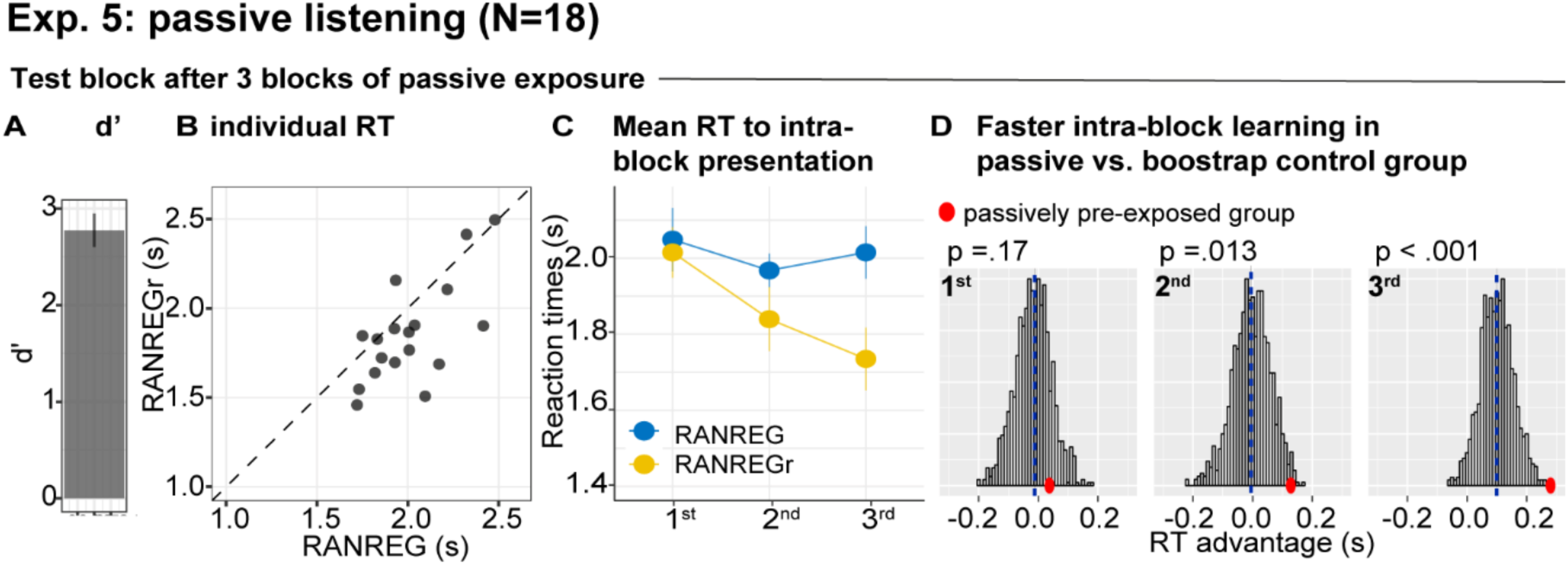
Experiment 5: implicit memory can form during passive exposure but does not immediately transfer to behaviour. During the 3 initial blocks, participants were asked to respond only to the STEP trials and ignore the other sounds. In the 4^th^ (test) block, they were instructed to also detect the RANREG transitions. **(A)** Sensitivity to emergence of regularity (d’) in the test block (mean d’ = 2.77 ± .73). Error bars indicate 1 s.e.m. **(B)** The relationship between RTs to the RANREG and RANREGr conditions in the test block. Each data point represents an individual participant. Dots below the diagonal indicate faster detection of RANREGr compared with RANREG. **(C)** RT to RANREG and RANREGr conditions separately for the 1^st^, 2^nd^ and 3^rd^ intra-block presentations. RT values for RANREG are determined by dividing the block duration into thirds and averaging across trials within each. Error bars indicate 1 s.e.m. **(D)** Bootstrap resampling-based distributions of the RT advantage for the 1^st^, 2^nd^ and 3^rd^ intra-block presentation from the control group (participants without previous passive exposure; see Methods). The mean of the distribution is indicated by blue dashed lines. Red dots indicate the data from the present experiment (passively pre-exposed group). One-tailed p-values are reported with each graph.

Following 3 blocks of passive exposure, the mean RT in the test block (Fig. 8 B) was faster in RANREGr than novel RANREG in the majority of participants. To understand the dynamics of the memory effect, we analysed the performance for each intra-block presentation of REGr (Fig. 8C-D). This was compared to the performance of a non pre-exposed ‘control’ group, formed by pooling block 1 data from several experiments (*Pooled data-block_1_*, see Methods). The plots in Fig. 8D show distributions of the RT advantage after the 1^st^, 2^nd^ and 3^rd^ REGr presentation in the control group (see Methods). The actual data from the test block in the present experiment are shown by the red dot. The RT advantage to the first presentation did not differ from the control group. However, a difference emerged after the 2^nd^ presentation. This suggests that by the second appearance of REGr in the ‘test’ block the passively pre-exposed group exhibited substantially better behaviour than that shown by non pre-exposed participants. The difference between the passively pre-exposed group and the control group grew further by the 3^rd^ presentation. Therefore, these results indicate that implicit memory was not present at the onset of the test block (as evidenced by the lack of an RT advantage), however learning occurred more rapidly in the pre-exposed listeners such that by the end of the test block, they exhibited a substantially higher RT advantage than that shown by the control group.

Explicit memory was poor (mean MCC = .064, see Fig. S2C) and did not correlate with the RT advantage measured in the test block [Spearman’s Rho = 0.235; p = 0.347].

### Across-experiment analysis reveals that most patterns are remembered and most participants exhibit implicit memory

We investigated the robustness of the memory effect for reoccurring patterns across different experiments. Fig. 9A shows the distribution of RTs for RANREG vs. RANREGr sequences pooled from block 3 data, (i.e., after 9 presentations of each REGr; approx. 25 minutes of listening) where most data from different experiments were available (the pilot experiment, Experiment 1A, 1B, 3, 4, S1, and S3). In Fig. 9B each dot represents the mean RT for RANREG vs. RANREGr sequences of an individual participant (N = 147). 88.4% of participants exhibited an RT advantage, which we interpret as revealing implicit memory for REGr.

**Fig. 9.**
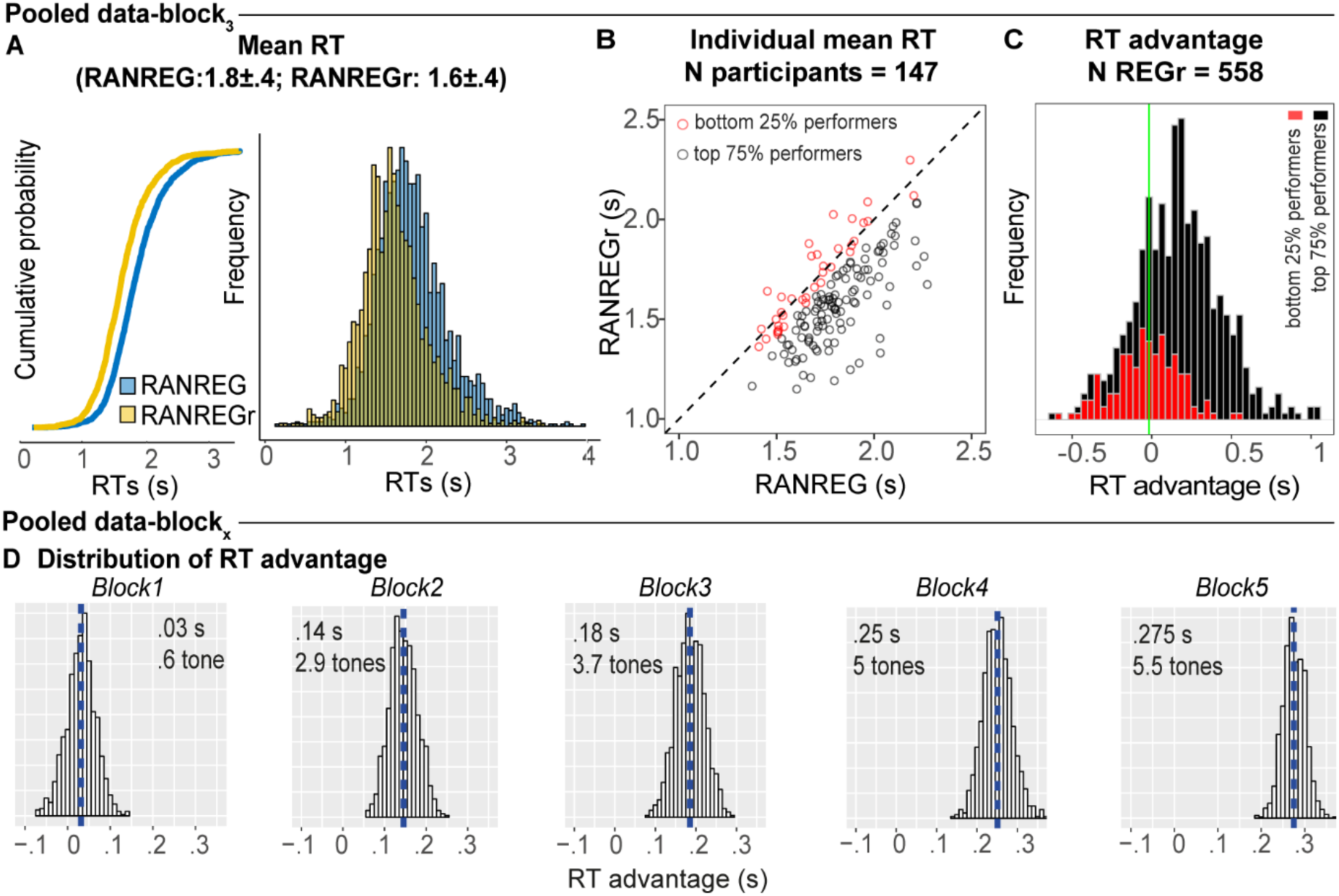
Individual variability in implicit memory. **(A)** Cumulative distribution function (left) and distribution (right) of RTs to RANREG and RANREGr pooled from block 3 of several experiments (see Methods). A two sample Kolmogorov-Smirnov test confirmed a significant difference in cumulative probability (D = 0.232, p-value < .001) **(B)** The relationship between RTs to the RANREG and RANREGr conditions in block 3. Each circle represents an individual participant. Participants in the bottom quartile (those who exhibited the smallest RT advantage) are shown in red. **(C)** Distribution of RT advantages across 558 different REGr patterns as measured after 3 blocks (9 reoccurrences). Values > 0 indicate faster RTs to REGr relative to novel REG. Stacked bars reflect the distribution across all REGr. REGr associated with the top 75% performers (as defined in A) are in black; those associated with the worst performers (bottom 25%) are in red. **(D)** Distributions of the RT advantage in each block. To estimate the distribution of the RT advantage across the population (of young, healthy participants) we pooled data from several experiments (see Methods) in which participants performed the standard regularity detection task. Pooled data-block_1_ reflects the distribution of RT advantage after one block (3 presentations of REGr), Pooled data-block_2_ reflects the distribution of the RT advantage after two blocks (6 presentations of REGr), etc. The distributions are computed via a bootstrapping process whereby on each iteration (1000 overall), data from 20 participants are chosen randomly (with replacement), to obtain an average RT advantage. The mean of each distribution is indicated by blue dashed lines. Overall these distributions demonstrate a robust emergence of an RT advantage after the first block.

We also tested the generality, across patterns, of the observed memory effect. It is important to note that all REGr were similar in the sense that all are composed from the same set of tones and only differed in the specific permutation of their order. Fig. 9C plots a distribution of the RT advantage per unique REGr (558 overall). Though the data are inherently noisy (RT is quantified as an average over only 3 presentations in block 3), 75.6 % of patterns exhibited a memory effect. Whilst amongst the low performing participants (lowest quartile; marked in red in Fig. 9B, C) only 48 % of the presented REGr were ‘remembered’ (as indicated by an RT advantage), the high performing participants learned 85.3% of REGr. This demonstrates that the observed effects are not driven by particularly ‘memorable’ REGr sequences. The same analysis run over block 5 data (not shown; # unique REGr = 165) showed that 91.5 % of REGr were associated with an RT advantage after 15 reoccurrences. Fig. 9D plots the distributions of group RT advantage per block, based on performance observed across all of the experiments reported (see Methods). A gradual build-up of RT advantage is seen across blocks reaching a mean of 5.5 tones by the end of block5.

Overall the results demonstrate that the memory effect generalizes to most (healthy, young) listeners and is not driven by particular memorable stimuli. These findings also inspire an interesting question about what cognitive abilities distinguish the low- from the high-performers.

## Discussion

We used rapid sequences of discrete sounds (Barascud et al., 2016; Southwell et al., 2017; Zhao et al., 2019) specifically structured to allow for detailed behavioural and model-based investigation of memory formation. All sequences were generated from a fixed set of 20 frequencies, with the only difference being the order in which these were presented. Participants performed a regularity detection task and were oblivious to the reoccurrences of certain regular patterns throughout the session. However, the pattern of reaction times to new vs. previously encountered regularities demonstrated that following limited exposure to reoccurring patterns listeners retained sequential information in long-term memory. This representation was implicit, resistant to interference, and preserved over remarkably long durations (over 7 weeks).

Statistics of pattern learning across experiments revealed that most patterns were remembered, and most participants exhibited a memory effect, although the size of this effect varied across individuals. Experimental results were successfully simulated by a memory-constrained model of sequential pattern acquisition suggesting that human memory may operate under similar constraints.

Overall the results establish the brain’s capacity to implicitly preserve sequential information in long-term memory and implicate an interplay between rapid and slow memory decay in supporting the formation of enduring memories of arbitrary sound sequences.

### Relationship to ‘noise memory’

The general behavioural pattern revealed here is reminiscent of the ‘noise memory’ effect first shown by Agus et al. (2010; see also Agus & Pressnitzer, 2013; Andrillon et al., 2015; Gold, Aizenman, Bond, & Sekuler, 2014; Keller & Sekuler, 2015; Luo, Tian, Song, Zhou, & Poeppel, 2013). In that study naïve listeners readily remembered reoccurring white-noise snippets presented amongst novel noise bursts. The learning was unsupervised, rapid, implicit and lasted upwards of 2 weeks.

Inspections of the nature of this memory revealed that it was robust to time reversal, and even to scrambling into bins as small as 10-20 ms (Agus, Thorpe, & Pressnitzer, 2010; Viswanathan, Rémy, Bacon-Macé, & Thorpe, 2016), indicating that the remembered features reflect local spectro-temporal idiosyncrasies within the reoccurring noise snippet. The apparent dependence of this memory on certain local features of the noise signal may also explain the high inter-sample variability often seen with this paradigm (i.e., the distinction between ‘memorable’ and ‘not memorable’ patterns; Agus et al., 2010; Viswanathan et al., 2016; Kang et al, 2017).

In contrast, here we focus on fast memory formation for *sequences* of discrete tones, distinguishable only by their specific order, and presented in a surrounding context of highly similar patterns (all sequences consisted of the same 20 ‘building blocks’). We showed that the vast majority of patterns were learned, revealing high sensitivity to reoccurring arbitrary frequency patterns despite the exceedingly rare reoccurrence rate (every ∼2.7 minutes; 5% of trials; in contrast to the much more frequent reoccurrence (< ∼15 seconds) in Agus et al (2010) and Kang et al (2017).

An important question for future work will be to determine whether these effects draw on similar or distinct neural systems. Agus et al. proposed that mechanisms based on spike-timing-dependent plasticity (STDP; Markram, Lübke, Frotscher, & Sakmann, 1997; Masquelier, Guyonneau, & Thorpe, 2008; Masquelier, Hugues, Deco, & Thorpe, 2009) may be possible neural underpinnings for rapid noise memory formation: repeatedly presented, but relatively temporally confined, spectro-temporal ‘constellations’ within the noise snippets may evoke coincident firing among auditory afferents leading to rapidly emerging selectivity for this feature in subsequent presentations of the same noise burst. Kang et al (2017) proposed that incorporating a degree of temporal integration can also account for similar effects observed with random click trains. It is possible that state-dependent neural place maps, as proposed by Kang et al (2017; see also Karmarkar & Buonomano, 2007; Lim, Lagoy, Shinn-Cunningham, & Gardner, 2017) incorporating an integration time of several hundred msec, may also underpin memory for discrete tone sequences. As will be discussed further below, the behavioural pattern reported in the present paper is consistent with sequential information being stored as short sub-sequences, i.e. without storing a representation of the full 20-item sequence.

### Relationship to ‘statistical learning’

A large body of work, broadly referred to as ‘statistical learning’ has demonstrated the brain’s capacity to discover repeating structure in random stimulus sequences (Conway & Christiansen, 2005; Frost, Armstrong, & Christiansen, 2019; Kim et al., 2009; Saffran, Johnson, Aslin, & Newport, 1999; Saffran & Kirkham, 2018). The classic paradigm (Saffran, Aslin, & Newport, 1996) involves a small ‘alphabet’ arranged into short ‘words’ (e.g., 3 syllables each). A few minutes’ exposure to such structured streams leads to learning of the statistical structure of the unfolding sequence such that subjects can distinguish the repeatedly occurring ‘words’ from a random arrangement of syllables.

Our results can be interpreted as reflecting similar processes. However, in contrast to the standard demonstrations which usually involve a small number of short words that are repeated many times, we show that a very sparse presentation of long patterns which are intermixed with many highly similar sequences, is sufficient for robust memories to be formed. In addition, the modelling suggests that listeners learn statistical dependencies of greater order than the first-order ‘transition probabilities’ usually considered in statistical learning research. The results also suggest that once learned, this statistical knowledge is subject to weakening over moderate timescales (within an experimental session) but subsequently remains intact in long-term memory.

More broadly, we demonstrate that the brain is tuned to discover and retain repeated structures in our acoustic environments, even when such reoccurrences are exceedingly infrequent and the signals are highly similar. Preserving as much information as possible from the unfolding sensory information is important for an organism as repetition in the environment often signals behaviourally relevant information and the relevance of events is not always immediately inferable (McDermott, Wrobleski, & Oxenham, 2011; Woods & McDermott, 2018). Our results hint at the heuristics utilized by the brain in determining how to reinforce retained representations of statistical structure in the sensory environment.

### What is being remembered?

We used reaction time (RT) as a proxy for memory formation. RT allowed us to determine how much information was required for listeners to detect repeating (REG) structure and to compare these measures with formal models of sequence processing. We hypothesized that reoccurrence would increase the weight of sequence components in memory resulting in faster detection of regularity. RT thus provided a sensitive means for tracking the formation and maintenance of such memories over time.

We observed that the RT to REGr steadily shortened with increasing number of reoccurrences, allowing us to measure the dynamics of memory trace establishment. The ‘RT advantage’, defined as the difference in RT between novel and reoccurring REG patterns, grew rapidly over the first 3 blocks (9 reoccurrences) and then stabilized, though evidence from Fig. 9D suggests a continuous slow growth throughout the experimental session. The absence of correlation between the familiarity test and the RT advantage suggests a dissociation between implicit memory and explicit recall abilities.

The RT advantage to REGr reflects an implicit memory of sequential structure (Exp. 1B). But what, specifically, is remembered? Clearly participants did not perfectly memorize the full pattern, in that this would have been associated with much faster RTs (e.g. as exhibited by the observer with unconstrained memory, Fig. 5A). Instead, by the end of block 3, the distribution of RT to REGr shifted leftwards by about 4 tones, without otherwise changing (Fig. 9A). Modelling suggests that this performance is consistent with a statistical-learning effect whereby the participants retained imperfect memory of patterns presented earlier in the experiment. These memories are not strong enough to prompt immediate recognition of a pattern heard in a past trial, but they are sufficiently strong to speed the recognition of that pattern once it begins repeating in the new trial.

The modelling (PPM) explicitly assumed that listeners represent the unfolding sequences in the form of n-grams. Previous computational, behavioural and neuroimaging studies (Barascud et al., 2016; Conklin & Witten, 1995; Egermann, Pearce, Wiggins, & McAdams, 2013; Pearce, Ruiz, Kapasi, Wiggins, & Bhattacharya, 2010; Pearce & Wiggins, 2004, 2006) demonstrated that PPM generalizes well to prediction of musical sequences and effectively accounts for psychological responses to melodies. In particular, PPM provided a good match to brain response latencies evoked by transitions between RAN and REG patterns (Barascud et al., 2016; Southwell & Chait, 2018), suggesting that listeners may rely on similar memory representations as those proposed by the model. Further support for this hypothesis is provided in Exp. 4, which demonstrated that the REGr RT advantage is robust to pattern phase shifts. This finding indicates that REG patterns are not encoded in memory as long rigid chunks of sequential items (Perruchet & Pacton, 2006; Thiessen, 2017), but instead represented as a collection of n-grams and their associated frequency counts, which allows for flexible retrieval. The more a listener is exposed to a given *n*-gram, the more salient this *n*-gram is in memory, leading to faster retrieval. We showed that listeners can learn at least 6 concurrently presented REGr patterns (Exp. 4 and Exp. S1). Important questions for future work involve understanding the capacity limits on this memory and the factors which might affect subsequent forgetting.

It is important to note that our listeners were placed in rather extreme conditions, both in terms of presentation rate of reoccurring targets and their complexity. We focused on implicit memory formation and therefore kept the reoccurrence of patterns deliberately sparse to avoid them becoming apparent to the participants. It is possible that relaxing that constraint would result in stronger (but perhaps explicit) memories. Additionally, all sequences shared the same long-term statistics (permutations of the same 20 tones) and hence high inter-pattern similarity. It is likely therefore that the memory weight difference between n-grams from novel vs. reoccurring patterns was small. Larger memory effects may be observed with more pronounced differences between sequences.

### Time scales of memory for sequences

The basic behavioural task required participants to detect the transitions from RAN to REG – namely the emergence of repeating structure. As such it fundamentally relied on auditory short-term memory: in order to detect REG patterns, the listener must compare incoming tones to those that occurred at least a cycle ago by relying on auditory short-term memory. The effect of reoccurrence also suggested that listeners draw on much longer-term memory whereby information about previously encountered sounds is retained over minutes (at least ∼ 2.7) between successive REGr presentations.

All of the reported experiments were subject to fixed presentation parameters (limited by practical issues related to providing breaks) whereby the experimental session was divided into blocks of roughly 8 minutes and REGr were presented 3 times per block. We therefore only have a relatively coarse estimate of the properties of the long-term memory store. Lengthening of inter-reoccurrence intervals was probed by introducing interrupting blocks where only novel patterns were presented (see Exp. S2A-B). Memory was largely maintained over roughly 10 minute intervals indicating a very slow long-term decay.

In conjunction, the results of Exp. 2 suggested that the short-term memory store is critical for pattern memory. Participants were markedly impaired at detecting pattern repetition when the two cycles were separated by a brief series of random tones (about 2 sec). Those conditions were also associated with substantially reduced long-term memories for the reoccurring patterns, suggesting that local reinforcement is critical for the formation of lasting memory traces.

These observations point to a critical interplay between a short (few seconds) and much longer (at least a few minutes) integration in the process of formation of robust, implicit long-term memories for reoccurring arbitrary sound sequences. Modelling suggested that memory decay parameters with short term memory half-life of 3.06 sec and long-term memory half-life of 500 sec provide a good account of the pattern of results exhibited by listeners.

Further work is required to understand how these decay parameters generalize across a range of experimental/sound settings and the neurobiological underpinnings of these effect. The hippocampus may play a key role in storing the statistics of unfolding sound sequences. Previous work has implicated the hippocampus in sensitivity to sensory patterns across rapid time scales (Aly, Ranganath, & Yonelinas, 2013; Stachenfeld, Botvinick, & Gershman, 2017; Yonelinas, 2013) and specifically in the process of discovering RAN-REG transitions (Barascud et al., 2016). There is also some evidence that hints at its possible role for supporting long-term memory for acoustic patterns (Kumar et al., 2014).

### Does sequence memory require attention?

The short-term memory mechanisms which allow listeners to discover the emergence of repeating structure (RANREG) in rapid tone sequences have been demonstrated to operate automatically, even when listeners’ attention is directed away from sound: brain activity recorded from naïve, distracted listeners reveals robust responses to RAN-REG transitions with latencies consistent with those expected from an ideal observer (Barascud et al, 2016; Southwell et al, 2017; Southwell & Chait, 2018).

In contrast, in Exp. 5 we demonstrated that the effect of longer-term statistical learning appears to require attentive processing in that there was no evidence for an immediate RT advantage when listeners moved from the passive exposure blocks to the active detection (‘test’) block. This suggests that the process of memory trace formation is not fully automatic, or does not immediately translate to behaviour. It is difficult to draw strong conclusions from the present experiment about the effect of attention per se. It is possible that the reduced memory effect in the passive condition was driven by issues related to decreased arousal or reward, known to modulate learning (Beste & Dinse, 2013; Braun, Wimmer, & Shohamy, 2018; Polley, Steinberg, & Merzenich, 2006; Yebra et al., 2019), and which likely distinguish active detection (where feedback was provided after each trial) from passive listening.

Importantly we showed that while implicit memory was not present at the onset of the test block, learning occurred more rapidly in the pre-exposed listeners, hinting at the presence of latent traces (Frankland, Josselyn, & Köhler, 2019) that contribute to faster instantiation of representations in memory once the sequences become behaviourally relevant.

## Methods

### Stimuli

Stimuli were sequences of 50-ms tone-pips of different frequencies generated at a sampling rate of 44.1 kHz and gated on and off with 5-ms raised cosine ramps. Twenty frequencies (logarithmically-spaced values between 222 and 2,000 Hz; 12% steps) were arranged in sequences with a total duration of 7 s (140 tones). The order in which these frequencies were successively distributed defined different conditions, that were otherwise identical in their spectral and timing profiles (see Fig. 1). RAN sequences consisted of tone-pips arranged in random order, with the constraint that adjacent tones were not of the same frequency. Each frequency occurred equiprobably across the sequence duration. The RANREG condition contained a transition between a random (RAN), and a regularly repeating pattern (REG): Sequences with initially randomly ordered tones changed into regularly repeating cycles of 20 frequencies (an overall cycle duration of 1000 ms; new on each trial). The change occurred between 3000 and 4000 ms after sequence onset such that each RANREG sequence contained between 3 to 4 REG cycles (only 2 in Exp. 2, see below). RAN and RANREG condition were generated anew for each trial and occurred equiprobably. Thus, each trial contained exactly the same frequency ‘building blocks’, with the same overall distribution, and only varied in the specific order of tone-pips.

Unbeknown to participants, a few different REG patterns (different for each participant) reoccurred identically several times within the session (RANREGr condition). Reoccurrences happened 3 times per block (every 2.7 minutes), and 9-15 times per session, depending on the number of blocks in the specific experiment. Note that the RAN part preceding each REGr was always novel. Reoccurrences were distributed within each block such that they occurred at the beginning (first third), middle and end of each block.

Two control conditions were also included within each block: sequences of tones of a fixed frequency (CONT), and sequences with a step change in frequency partway through the trial (STEP). The STEP condition served as a measure of individuals’ reaction time to simple acoustic changes. The RT to STEP was subtracted from the RT to RANREG sequences to obtain a lower bound measure of computation time required to detect the transition. The inter-stimulus interval was jittered between 1400 and 1800 ms.

Participants were instructed to respond, by pressing a keyboard button, as soon as possible after detecting a RANREG transition. Feedback about response accuracy and speed was delivered at the end of each trial. Since RT is a key performance measure in these experiments, it was important to motivate the participants to respond as quickly as possible. The feedback was given based on our previous work (Barascud et al., 2016), and consisted of a green circle if the response fell within 2200 ms from the regularity onset in the RANREG conditions, or within 300 ms from the change of tone in the STEP condition. For slower RTs, an orange circle (between 2200 – 2600 ms in the RANREG conditions, and between 300 – 600 ms in the STEP condition) or a red circle were displayed. It was explained to participants that they should strive to obtain as much ‘green’ or ‘orange’ feedback as possible. The experimental session was delivered in ∼8 min blocks, separated by brief breaks. Stimuli were presented with PsychToolBox in MATLAB (9.2.0, R2017a) in an acoustically shielded room and at a comfortable listening level (self-adjusted by each listener).

### Participant numbers

We initially ran a pilot experiment (N=20, 16 females, age 23.5 ± 2.95 years) which consisted of five consecutive blocks as in Exp. 1A. The effect size for the main effect of condition (RANREG vs. RANREGr) was η_p_^2^ = .48 and η_p_^2^ = .79 after the first 3 and 5 blocks respectively. Using η_p_^2^ =0.48 for a prospective power calculation yielded a required sample size of N = 11. We decided to increase our sample size up to N = 20 to account for possible drop outs due to low accuracy. The research ethics committee of University College London approved the experiment, and written informed consent was obtained from each participant.

### Experiment 1A

The transition detection task was performed in three sessions: five blocks on day one (‘day1’), one block after 24 hours (‘24h’) and another block after 7 weeks (‘7w’). Each block consisted of 60 stimuli (∼8 minutes duration; 3 RANREGr × 3 reoccurrences per block, 18 RANREG, 27 RAN, 3 STEP, and 3 CONT). Before starting, a short training block of 12 trials (with the same conditions as in the main experiment, but no RANREGr) was performed to acquaint participants with the task.

The **familiarity task** was performed at the end of each session (day1, 24h, 7w). In these tests the three REGr patterns were randomly intermixed with 18 novel REG sequences. Participants were informed that a ‘handful’ of patterns reoccurred during the just completed session and asked to indicate, by means of a button press, if each presented pattern sounded ‘familiar’. Feedback about the response accuracy and speed was delivered after each trial.

#### Participants

Twenty paid individuals (ten females; average age, 24.4 ± 3.03 years) took part in the experiment. Because of poor accuracy (d’ < 2 after the first block), one participant was excluded from the analysis. We were able to test only 14 participants after 7 weeks (8 females; average age, 24.7 ± 3.02 years). No participant reported hearing difficulties.

### Experiment 1B

Participants performed the transition detection task for 6 consecutive blocks consisting of the same set of stimuli described for Exp. 1A. In the 5^th^ block, each REGr was time reversed.

#### Participants

Twenty paid individuals (13 females; average age = 24.25 ± 3.58 years) took part in the experiment. No participant reported hearing difficulties.

### Experiment 2

The stimulus set in the initial 4 blocks contained RANREG and RANREGr stimuli, as before, except they contained only 2 repeating cycles after the transition. To understand whether immediate repetition is necessary for memory to be formed two further conditions were used: PATinRAN stimuli contained two identical 20 tone patterns embedded amongst random tones (mean separation of 1.7 s; drawn randomly from a range .5 – 2.9; the second appearance always occurred at the end of the sequence as shown in Fig. 3A). Similar to REGr, 3 different PAT were designated as reoccurring across trials (different for each participant; 3 reoccurrences per block). The embedding RAN sequence and the spacing between the two PAT patterns were randomly set for each reoccurrence. Overall each block contained 82 stimuli (36 RAN, 9 RANREG, 9 RANREGr, 9 PATinRAN, 9 PATinRANr, 5 STEP, 5 CONT), with ISI between 2.4 and 2.8 s. Reoccurrences of RANREGr and PATinRANr occurred approximately every 3.6 minutes.

Participants were informed of the presence of PATinRAN and RANREG stimuli (but were naïve about RANREGr and PATinRANr) and were instructed to indicate, by button press, if they detected the presence of a repeating pattern with the just-heard sequence. Feedback was provided at the end of each trial as in the above experiments, except that in the PATinRAN conditions we delivered a green circle if the response fell within 1200 ms from the second cycle onset, a red circle if the response was slower that 1600 ms, and an orange one if it fell in between. It was explained to participants that they should be fast but prioritise accuracy, given the generally difficult level of the task.

In order to quantify any memory effects, in the 5^th^ block (‘test’ block) each of PATinRANr sequences were replaced by sequences with the 2 cycles set adjacently. We will refer to this condition as RANREGr*. The test block contained 36 RAN, 18 RANREG, 9 RANREGr, 9 RANREGr*, 5 STEP, 5 CONT Stimuli were about 5.45 ms long (∼ 109 tones).

#### Participants

Given the task complexity and expectation for a reduced SNR, we increased the number of participants, a-priori, by 50% relative to the previous experiment. Thirty paid individuals (twenty females; average age, 24.26 ± 3.8 years) took part in the experiment. No participants reported hearing difficulties.

### Experiment 3

This experiment consisted of two days of testing. On the first day participants performed a transition detection task as in Exp. 1A, but two different sets of reoccurring patterns (REGr1 and REGr2; 3 different patterns each) were presented. RANREGr1 was presented over the first 3 blocks, and RANREGr2 over the subsequent 3 blocks. On day two (after 24 hours) participants returned to the lab to perform two test blocks for the two sets of reoccurring regularities, REGr1 and REGr2 (order counterbalanced across participants).

#### Participants

We initially ran 20 participants (1 excluded from analysis), but decided to run an additional 10 participants (+2 excluded), to increase the SNR for the memory effects observed for RANREGr1 and RANREGr2 conditions on day two. The results with N =19 yielded qualitatively similar results (see Fig. S3). Thirty-two paid individuals (twenty females; average age, 24.5 ± 3.8 years) took part in the experiment. No participant reported hearing difficulties. Because of poor accuracy (d’ < 2 after the first block), three participants were excluded from the analysis.

### Experiment 4

Participants performed the detection task through four consecutive blocks of 82 stimuli each. The stimulus set included the same conditions as described for Exp. 1A, but with six, instead of three, REGr sequences, each presented three times within a block (6 RANREGr × 3 reoccurrences per block, 18 RANREG, 36 RAN, 5 STEP, and 5 CONT). In block 4, REGr were phase shifted (see examples in Fig. 7A). To ensure uniform sampling of possible phase shifts, for each REGr in block 4, each of the three intra-block presentations was subject to pattern phase shift of 2 to 7, 8 to 13, or 14 to 19 tones from the onset of the original pattern. The phase shift was determined independently for each REGr and each intra-block presentation. Stimulus duration was 6.5 s, and the transition time was between 3 and 3.5 s from the sequence onset. Different REGr patterns reoccurred sparsely (every ∼3.4 minutes) across trials and blocks.

#### Participants

Twenty paid individuals (fourteen females; average age, 23.5 ± 3.2 years) took part in the experiment. No participant reported hearing difficulties.

### Experiment 5

The experiment consisted of 4 blocks. The stimulus structure was as in Exp. 1A, except that for the first 3 blocks participants were instructed to respond to STEP changes only. They received no explanation about the regularity structure of the stimuli, and performed no practice. On the fourth block, they were instructed to detect RANREG transitions in addition to STEP transitions. Each block contained 72 stimuli (3 RANREGr × 3 reoccurrences per block, 18 RANREG, 27 RAN, 9 STEP, and 9 CONT; ISI between 900 and 1300); the number of STEP and CONT trials was increased relative to that in Experiment 1A due to the task change. As in Exp. 1A, participants performed the familiarity task at the end of the session.

#### Participants

Nineteen paid individuals (14 females; average age, 23.4 ± 3.1 years) took part in the experiment. No participant reported hearing difficulties. Because of poor accuracy (d’ < 1), one participant was excluded from the analysis.

### Statistical analysis

In the transition detection task, two indexes of performance were computed: sensitivity (d’) and reaction times (RTs).

For each participant and each block, d’ was quantified over trials (collapsed over RANREG and RANREGr) to give a general measure of sensitivity to the presence of regularities. Participants who showed d’ < 2 after the first block of the transition detection task were excluded from the analysis. Because Exp. 5 had only one ‘active’ block and no previous training, we adopted a more lenient exclusion criterion of d’ <1. Note that d’ was not available in Exp. 2 because of the intermixed nature of the presentation of RANREG and PATinRAN stimuli. To quantify performance, we therefore focus on hit rates and false alarms. For the purpose of participant exclusion, we computed an overall d’ (collapsing across conditions) and set the threshold at d’ < 1.5.

Only RTs of correct trials (hits) were analysed. In all experiments, RT was defined as the time difference between the onset of the regular pattern (‘nominal transition’ in Fig. 1) and the participant’s button press. However, Exp. 2 contained conditions with non-contiguous pattern presentations. RT was therefore computed from the onset of the second cycle (as indicated in Fig. 3A). Across all experiments, RTs which occurred before the regularity was physically detectable – before the ‘effective transition’ (see Fig. 1; ∼ 1.3% of the trials) were considered to indicate a false positive and excluded from the analysis. To control for individual latency of motor response to a simple acoustic change, RTs were then ‘baselined’ by subtracting the individual mean RT to the STEP transition. Moreover, for each participant and block, the RTs beyond 2 SD from the mean were discarded.

To quantify the formation of a memory trace over REGr presentations, RT were averaged to yield a mean RT per condition per subject per block. Therefore, RT to RANREGr were based on nine trials (3 REGr × 3 presentations per block). However, to evaluate the immediate presence of a memory trace following certain experimental manipulations (e.g. in Experiments 2 and 5) or when re-testing after 24 hours or 7 weeks (as in Exp. 1) we also analysed RT for each intra-block presentation (the first, second and third intra-block instance of a REGr pattern; see Fig. S1). To calculate the ‘RT advantage’ for each intra-block presentation, mean RTs of 1^st^, 2^nd^ or 3^rd^ intra-block presentation (averaged across the different REGr) were subtracted from the mean RTs of REG which occurred at the beginning (first third), middle or end of each block.

Performance was statistically tested with linear analyses of variance (ANOVA) implemented in the R environment (version 0.99.320) using the ‘ezANOVA’ function (Michael Lawrence, 2016). The analysis of d’ modelled the repeated measures factor block (1: N blocks). The analysis on RTs modelled the repeated measures factors: condition (RANREG/RANREGr), block (1: N blocks), and their interaction term. P-values were Greenhouse-Geisser adjusted when sphericity assumptions were violated. Post hoc t-tests were used to test for differences in performance between conditions across blocks, or experiments, and Bonferroni correction was applied based on the number of the comparisons.

As a benchmark (see Fig. 9D) across which to compare the effect of various manipulations on the RT advantage (e.g. Fig. 8D, S5C-G), we pooled data from several experiments to obtain a distribution of RT advantage values after each block: *Pooled data-block_1_*, *Pooled data-block_2_*, *Pooled data-block_3_* were formed by pooling block 1, 2 or 3 data, respectively, from Experiments 1A, 1B, 3, 4, S1, S3, and pilot experiment identical to Exp. 1 (total N=147). *Pooled data-block_4_* was formed by pooling block 4 data from Experiments 1A, 1B, S1, S3 and the pilot (total N=98), and *Pooled data-block_5_* by pooling block 5 from Experiments 1A, S1 and the pilot (total N=58). To obtain distributions of expected RT advantage values, data in each set were subjected to bootstrap resampling (1000 iterations) where, on each iteration, N random participants (N= number of participants in the experiment under examination) were drawn from the full pool, and their mean RT advantage (RANREG-RANREGr) was computed. This procedure yielded a distribution to which the actual data from the experiment under examination were compared. The p values provided (e.g. Fig. 8D, S5C-G) reflect the probability of the measured RT advantage (red dots in the relevant figures) relative to the benchmark distribution.

### Analysis of the familiarity task

The familiarity measurement required participants to categorize the presented patterns into ‘familiar’ (REGr) or ‘new’ (REG). Each REGr was presented once only, to avoid learning during the testing session and hence the ‘familiar’ category included only 3 items (6 in Exp. S1). These were presented among a larger set of foils (18 in Exp. 1A and Exp. 5, 36 in Exp. S1). Due to the small number of REGr, standard signal detection approaches are not useable. Instead we computed the MCC score, which is a measure of the quality of a binary classification, applicable even when classes are of different sizes (Boughorbel, Jarray, & El-Anbari, 2017; Powers, 2007). The coefficient ranges between 1 (perfect classification) to −1 (total misclassification) and is calculated using the following formula: 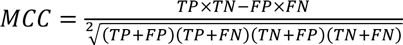. Where TP=number of true positives; TN=number of true negatives; FP=number of false positives; FN=number of false negatives. The MCC scores obtained for each participant in Exp. 1A, S1, and 5 are reported in Fig. S3.

### PPM-decay Model

Prediction by Partial Matching (PPM) is a Markov modelling technique (Cleary & Witten, 1984) that models statistical structure within symbolic sequences by tabulating occurrences of *n*-grams within a training dataset. PPM is a variable-order Markov model, meaning that it generates predictions by combining *n*-gram models of different orders; here we use a model combination technique called ‘interpolated smoothing’ (Bunton 1996, 1997; see also Pearce & Wiggins, 2004; Harrison et al., 2020; for more details). This approach combines the advantages of both the structural specificity afforded by high-order n-gram predictions and the statistical reliability afforded by low-order n-gram predictions.

The PPM models used in prior cognitive research (Barascud et al., 2016; Cheung et al., 2019; Gold, Pearce, Mas-Herrero, Dagher, & Zatorre, 2019) have a ‘perfect’ memory, in that historic n-gram observations are preserved with the same fidelity as recent events, and are weighted the same in prediction generation. Noting that human memory exhibits clear capacity limitations and recency effects, Harrison et al. (2020) modified PPM to incorporate a customizable decay kernel, whereby historic n-gram observations are down-weighted as a function of the time elapsed and the consequent n-grams observed since the initial observation. Modelling reaction-time data from a RANREG paradigm similar to Barascud et al. (2016), Harrison et al. concluded in favour of a capacity-limited high-fidelity echoic memory buffer followed by a lower-fidelity short-term memory phase with exponential decay. We likewise use an echoic-memory phase and a short-term memory phase in the present work, but add a slower-decaying long-term memory phase in order to capture the long-term learning observed in the present experiment.

The modelling aimed to reproduce behavioural performance qualitatively rather quantitatively. Many simplifications are made including That Inter-sequence intervals, and breaks between experimental blocks are modelled at a fixed rate of 1 sec. We explored various parameter settings for the model, and retained the configuration that best reproduced the observed behavioural patterns in Experiments 1A, 2, and S2A (Fig. 5, and Fig. S5D), which represent the key manipulations of memory duration. The resulting parameters are listed in Table 1; the decay kernel is plotted in Fig. 4A. Further implementation details are described in Harrison et al. (2020). The model outputs a conditional probability estimate for each tone in each sequence experienced throughout an experiment, which we convert to information content (the negative log probability in base 2). An implementation of this model is freely available in our open-source R package ‘ppm’ (https://github.com/pmcharrison/ppm).

To identify changes in the information content profile corresponding to the RANDREG transition on a given trial, we use the nonparametric changepoint detection algorithm of Ross, Tasoulis, & Adams (2011), which sequentially applies the Mann-Whitney test to identify changes in a time series’ location while controlling for Type I error. Here the target Type I error rate was set to 1 in 10,000 tones. Note that, for simplicity, the change point detection algorithm is free of memory constraints. Human listeners likely use a rougher (less detailed) statistical representation for transition detection.

## Supporting information

sounds

## Acknowledgments

We thank Barathy Ganeshakumara for help with the behavioural data collection and Alain de Cheveigne for comments and discussion. This research was supported by a BBSRC grant (BB/P003745/1) to MC.

## Competing interests

On behalf of all authors, the corresponding author declares no financial and non-financial competing interests.

## Data Availability Statement

The datasets for this study can be found in the OSF repository (https://osf.io/dtzs3/; DOI 10.17605/OSF.IO/DTZS3).

## Supplementary Media

See ‘sounds.zip’ folder in the OSF repository (https://osf.io/dtzs3/; DOI 10.17605/OSF.IO/DTZS3). This contains examples of RAN and RANREG stimuli as used in the reported experiments.

---

**Fig. S1.**
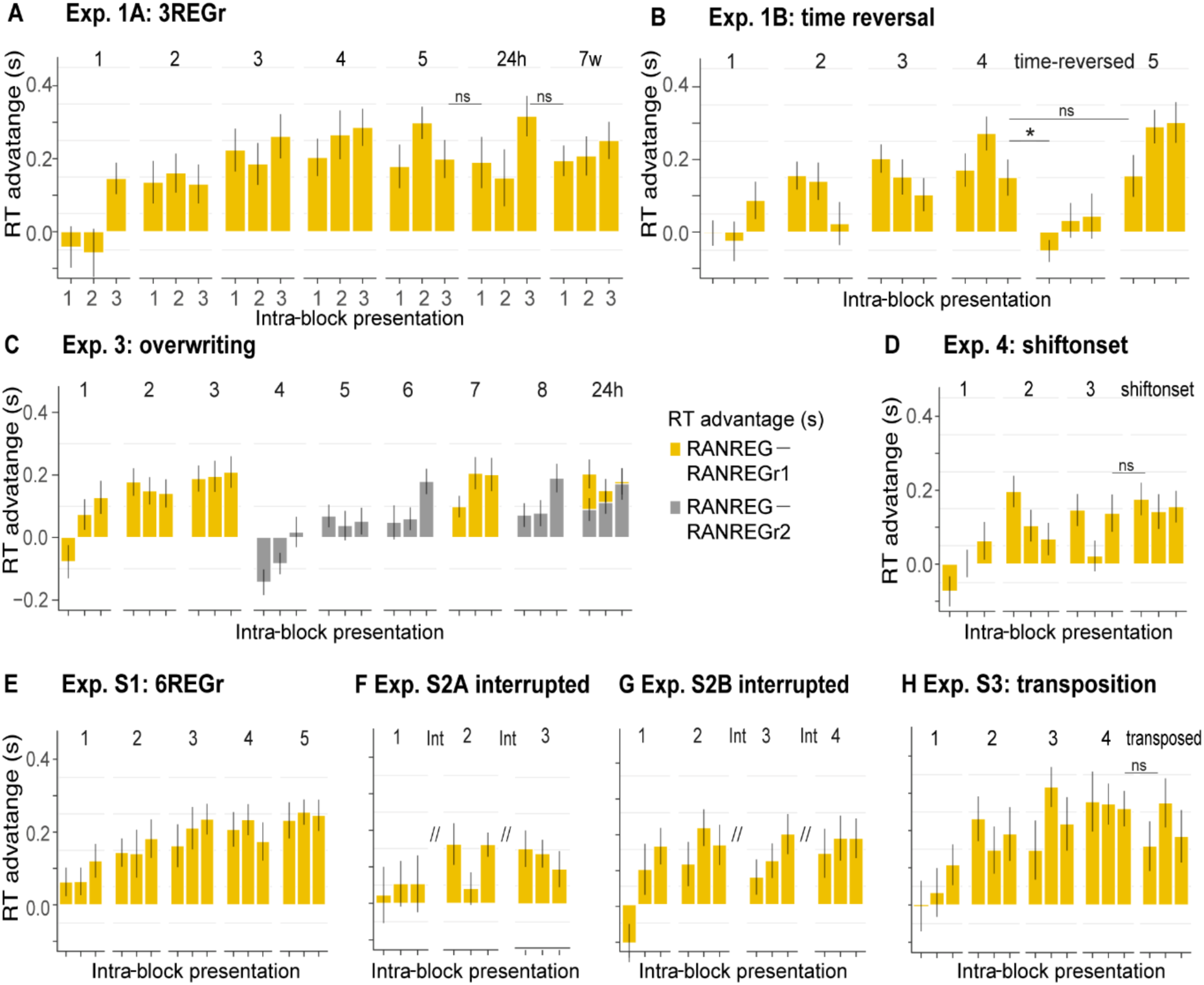
RT advantage for each intra-block presentation. These plots, from each of the reported experiments, depict the progressive emergence of an RT advantage with each presentation of REGr. Plotted values correspond to the RT advantage of REGr for each intra-block presentation (blocks are indicated on the top row of each graph). RTs of 1^st^, 2^nd^ or 3^rd^ intra-block presentations were averaged across the different REGr (6 in Exp S1; 3 in all the other experiments), and RTs to novel REG were averaged across trials which occurred at the beginning (first third), middle or end of each block. Error bars indicate 1 s.e.m. Note that the RT for REGr is computed based on 3 (or 6) trials and the effects are therefore rather noisy **(A)** Experiment 1A. There was no significant difference between the last presentation in block 5, and the first presentation after 24 hours, or between the last presentation after 24hours and the first presentation after 7 weeks, indicating that the formed memory trace was preserved long term **(B)** Experiment 1B. The RT advantage drops when REGr are time reversed and is restored when the original REGr are re-introduced (block 5) **(C)** Experiment 3. Dynamics of RT. The RT advantage for a set of reoccurring patterns (REG1; yellow traces) is not affected by the presentation of another set of REGr (REGr2) in blocks 4-6. **(D)** Experiment 4. The RT advantage is preserved after the introduction of a REGr phase shift. **(E)** Experiment S1. A progressive RT advantage emerges even when 6 different REGr are presented. **(F)** Experiment S2A. RT advantage is preserved over ‘interrupting’ blocks (see supplementary experiments below) **(G)** Experiment S2B. RT advantage is preserved over ‘interrupting’ blocks (see supplementary experiments below) **(H)** Experiment S3. The RT advantage is preserved following frequency transposition of the REGr pattern (see supplementary experiments below).

**Fig. S2.**
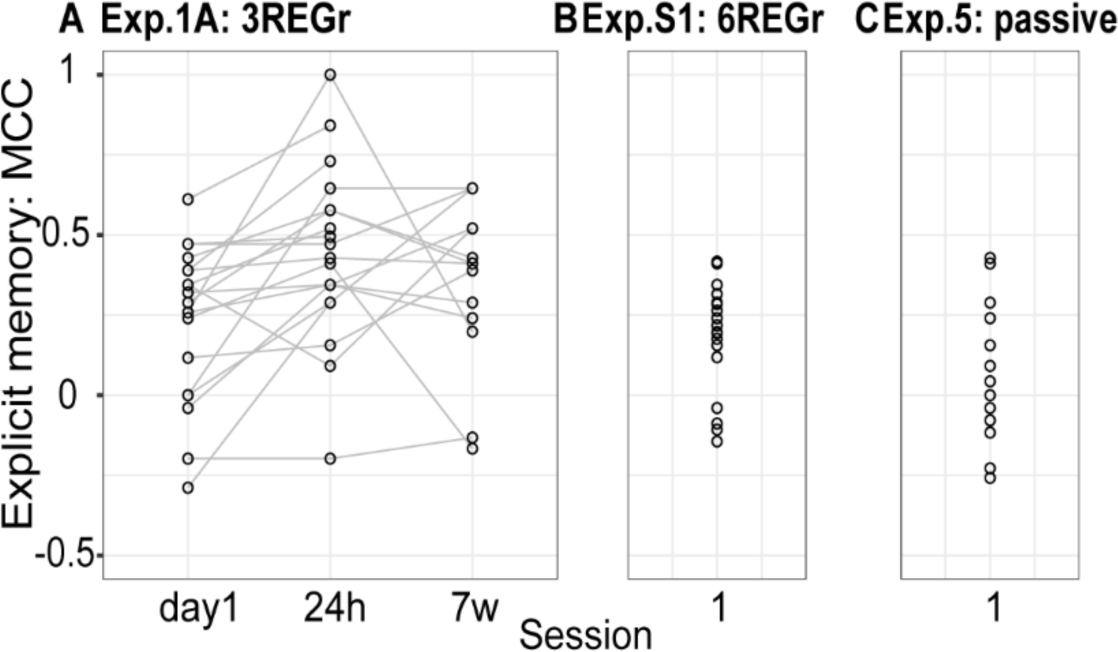
Explicit recognition estimates. MCC coefficient (refer to Methods) computed for the familiarity task performed after the regularity detection task in (A) Exp. 1A, (B) Exp. S1, and (C) Exp. 5. Each dot represents an individual participant. MCC was low overall, indicating low explicit recognition and did not correlate with the RT advantage (refer to main text).

**Fig. S3.**
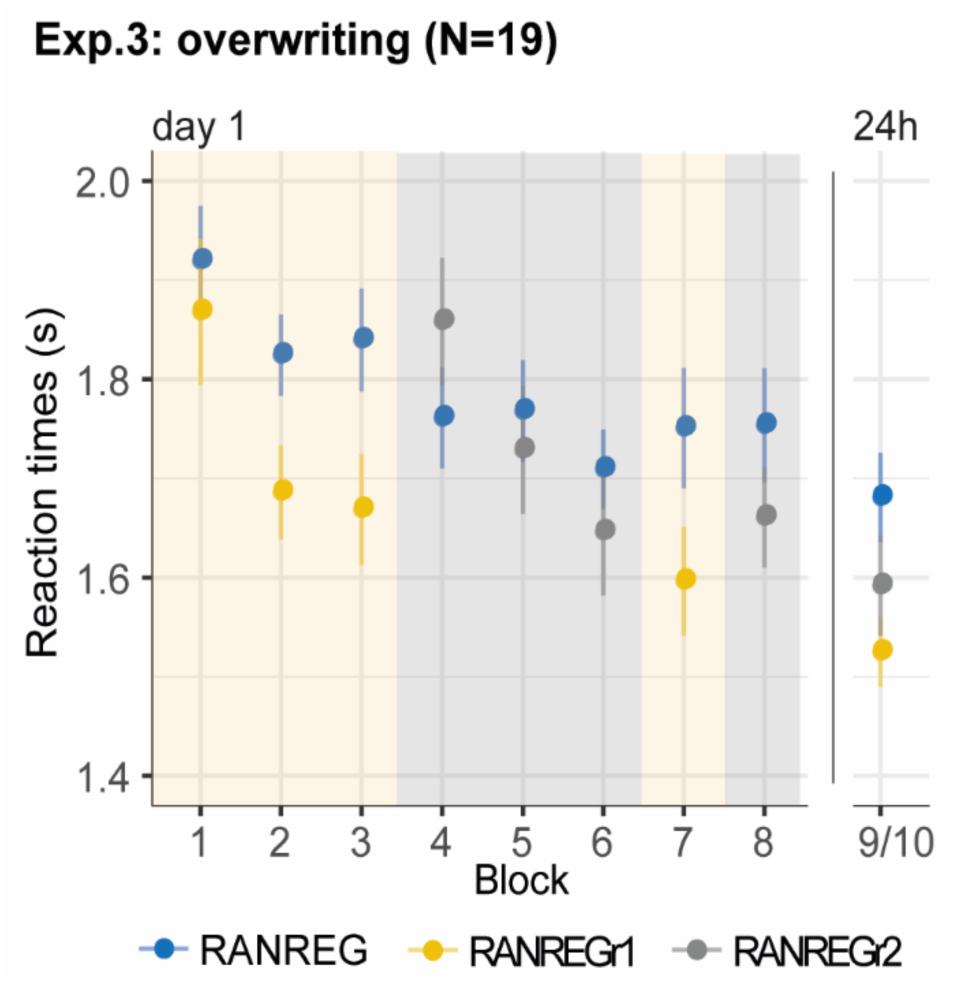
Experiment 3 (N=19). The overall pattern is identical to that observed with N=30 participants (reported in the main text). The RT advantage for the first set of REGr1 observed across the first 3 blocks [main effect of condition: F(1,18) = 24.16, p <.001, η_p_^2^ =.57; main effect of block: F(3,54) = 11.47, p < .001, η_p_^2^ =.39; condition by block interaction: F(3, 54) = 3.08, p = .035, η_p_^2^ =.15] was not perturbed after the presentation of the second set of reoccurring sequences [RT advantage for RANREG1 in block 3 vs. block 7: t(18) = .403, p =.691].

## Supplementary Experiments

### Experiment S1: Implicit memory for 6 concurrent patterns

In this experiment (Fig. S4) we probed implicit memory capacity by doubling the number of regularities to be memorised (6 different REGr per participant).

#### Methods

The transition detection task was identical to Exp. 1A, but 6 different REGr were presented per participant. Similar to Exp. 1A, participants performed the familiarity task after the transition detection task, in which the 6 REGr trials were randomly intermixed with 36 novel REG sequences.

##### Participants

Twenty paid individuals (seventeen females; average age, 24.5 ± 3.8 years) took part in the study. No participant reported hearing difficulties. Because of poor accuracy (d’ < 2 after the first block), one participant was excluded from the analysis.

#### Results

Overall, the same pattern of performance as in Exp. 1A was demonstrated.

Fig. S4-B reveals a progressively larger RT advantage for RANREGr [main effect of condition: F(1,18) = 71.76, p < .001, η_p_^2^ = .80; main effect of block: F(4, 72) = 4.19, p =.045, η_p_^2^ = .22; interaction condition by block: F(4,72) = 7.26, p < .001, η_p_^2^ = .29]. A significantly faster response (80 ms; 1.6 tones) for RANREGr relative to RANREG was observed already by the end of the first block [t(18) = 3.512, p =.012]. It grew across the following blocks (all ps < .001), and reached 244 ms (4.9 tones) in the fifth block, consistent with Exp. 1A [RT advantage in Exp 1A vs. Exp.2: independent sample t(36) = .515, p =.609].

Fig. S4-C shows the mean RT advantage for RANREGr for each individual in block 5. Implicit memory was exhibited by all participants by the end of the session.

Explicit memory (probed in the same way as described for Exp. 1A) was poor (mean MCC =.178, Fig. S2-B) and did not correlate with the RT advantage in block 5 (spearman’s Rho=0.091; p=0.710).

**Fig. S4.**
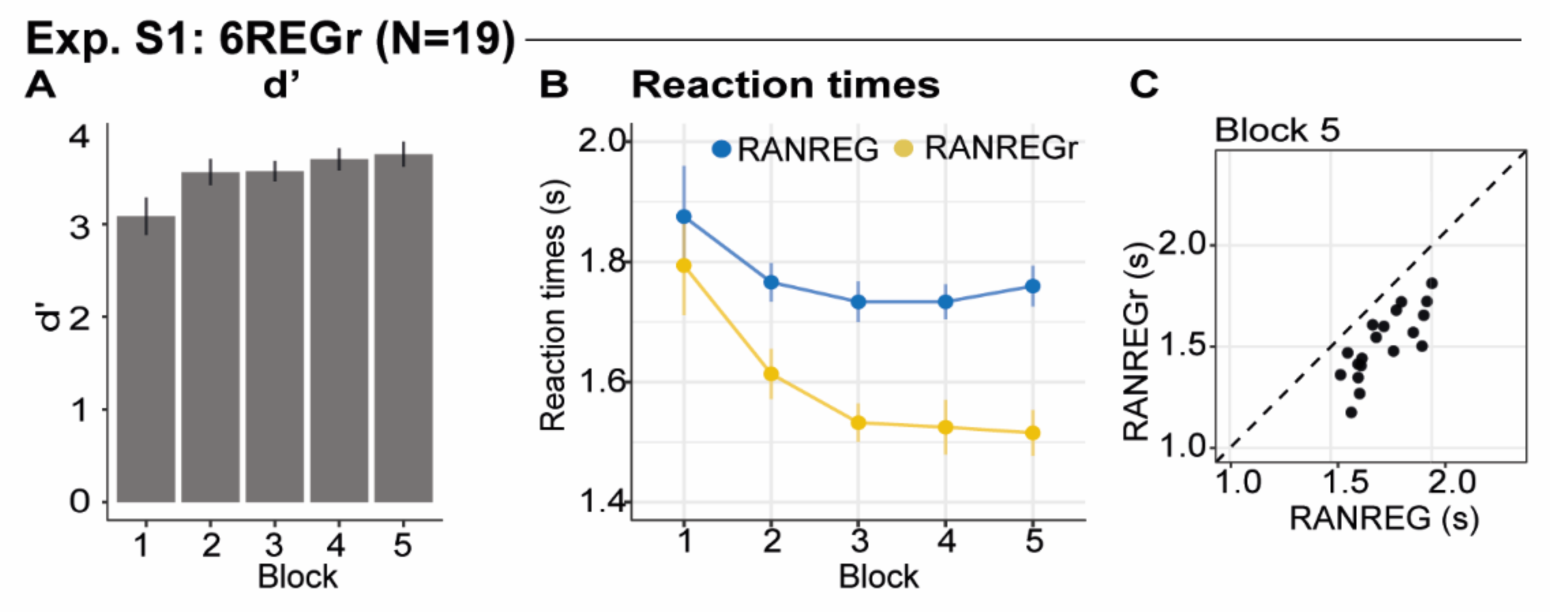
Experiment S1: implicit memory for 6 concurrent patterns. **(A)** Sensitivity to emergence of regularity (d’) across blocks. **(B)** RT to the RAN to REG transition in RANREG and RANREGr conditions across blocks. **(C)** The relationship between RTs to the RANREG and RANREGr conditions in block 5. Each dot represents an individual participant. All participants exhibited implicit memory of reoccurring patterns by the end of the 5^th^ block.

### Experiment S2A, B: The memory trace is weakened, but not abolished by interrupting blocks

Although reoccurrence of regularities was quite sparse in Exp. 1A (every ∼ 2.7 minutes), they were presented regularly over 5 blocks. Here, we asked whether memory formation can be interrupted by introducing a delay of 10 minutes (‘interrupting blocks’ in which REGr were not presented) between ‘standard blocks’.

#### Methods

These experiments involved the same transition detection task as in Exp. 1A, but ‘interrupting blocks’, in which RANREGr condition was not presented, were introduced between ‘standard blocks’. The ‘interrupting blocks’ were block 2 and 4 in experiment S2A, block 3 and 5 in experiment S2B. Across 5 blocks, in experiment S2A participants were presented with 27 RANREGr, 108 RANREG, 135 RAN, 15 STEP, and 15 CONT. Across 6 blocks, in experiment S2B participants were presented with 36 RANREGr, 126 RANREG, 162 RAN, 18 STEP, and 18 CONT.

##### Participants of experiment S2A

Nineteen paid individuals (13 females; average age, 23.8 ± 4.7 years) took part in the study. No participant reported hearing difficulties. Because of poor accuracy (d’ < 2 after the first block), one participant was excluded from the analysis.

##### Participants of experiment S2B

Twenty paid individuals (10 females; average age, 23.8 ± 4.00 years) took part in the study. No participant reported hearing difficulties. Because of poor accuracy (d’ < 2 after the first block), one participant was excluded from the analysis.

#### Results

In Exp. S2A, an interrupting block was inserted after each standard block (Fig. S5-B). The RT data demonstrated a RT advantage to reoccurring vs. novel regularities (∼130 ms – 2.6 tones by the end of the third standard block), which did not improve substantially across blocks [main effect of condition: F(1,17) = 35.03, p < .001, η_p_^2^ = .67; main effect of block: F(2, 34) = 10.67, p < .001, η_p_^2^ = .39; no interaction: F(2, 34) = 3.03, p = .061, η_p_^2^ = .15]. The RT advantage here was smaller than that typically observed after 3 consecutive blocks (∼180 ms – 3.7 tones in *Pooled data-block_3_*; Fig. S5-C; difference significant at p=0.027 based on bootstrap resampling; see Methods).

In Experiment S2B, we introduced the first interrupting block after block 2 in order to allow for the memory trace to emerge (see Fig. S5-F). The RT advantage in the 2^nd^ block was similar to that observed in the control (*Pooled data-block_2_*: p =0.48), but no considerable improvement was observed across blocks thereafter [main effect of condition: F(1,18) = 74.93, p < .001, η_p_^2^ = .81; main effect of block: F(3, 54) = 11.19, p < .001, η_p_^2^ = .38; no interaction: F(2, 54) = 2.56, p = .064 η_p_^2^ = .12]. The RT advantage in the blocks thereafter was indeed smaller than under ‘uninterrupted’ control conditions (block 3 vs. *Pooled data-block_3_*: p =.071; block 4 vs. *Pooled data-block_4_* : p =.013, see Fig. S5G).

These results suggest that the memory trace for REGr can withstand quite substantial interruptions: suspending the regular reoccurrences of REGr (by introducing ‘interrupting blocks’) resulted in a largely maintained memory, though there was evidence for a somewhat stagnated RT advantage.

### Modelling Exp. S2A

The performance of the unconstrained PPM model (Fig. S2-D, top), was not affected by the interruptions (also compare this Fig. with Fig5-A in the main text) In contrast, in the memory-decay PPM model inserting ‘interrupting’ blocks has the effect of reducing the memory traces of previously heard regularities. The constrained model shows somewhat worse performance relative to the constrained model in Exp. 1A, consistent with human effects.

**Fig. S5.**
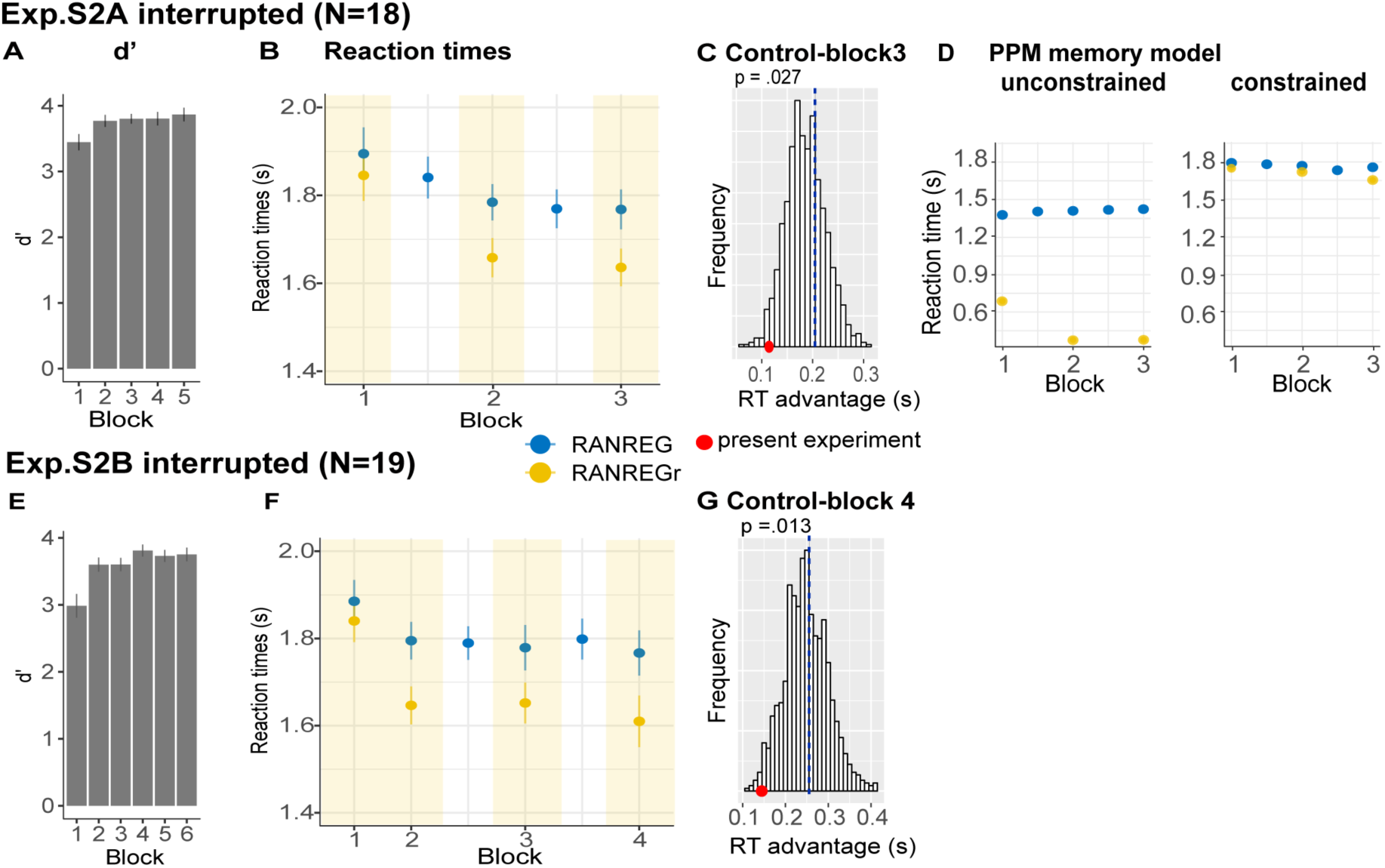
Experiment S2A and S2B: the memory trace is weakened, but not abolished, by interrupting blocks. **(A-D) Exp. S2A: (A)** Sensitivity to emergence of regularity (d’) across blocks in experiment S2A. Error bars indicate 1 s.e.m. **(B)** RTs to transition in RANREG and RANREGr across blocks. Error bars indicate 1 s.e.m. Yellow shading indicates blocks where REGr were present. **(C)** Bootstrap resampling-based distributions of RT advantage after 3 uninterrupted blocks (Pooled data-block3; see Methods). The red dot indicates the RT advantage measured after block 3 in the present experiment. **(D)** Unconstrained vs. Constrained memory results. Error bars indicate 1 s.e.m. **(E-F) Exp. S2B: (F)** Sensitivity to emergence of regularity (d’) across blocks for experiment S2B Error bars indicate 1 s.e.m. **(F)** RTs to the transition in RANREG and RANREGr across blocks. Error bars indicate 1 s.e.m. Yellow shading indicates blocks where REGr were present. **(G)** Bootstrap resampling-based distributions of RT advantage after 4^th^ blocks (Pooled data-block4; see methods). The red dot indicates the RT advantage measured after block 4 in the present experiment.

### Experiment S3: Implicit memory is robust to pattern transposition

We tested whether the implicit memory for reoccurring sequences generalises to versions in which relative relationships within the stimulus (pitch intervals) are preserved, while absolute information (the frequency values themselves) are manipulated.

#### Methods

The stimulus set included the same conditions as described for Exp. 1A, but with the following differences: RAN sequences were generated from a pool of twenty-six frequencies (logarithmically-spaced values between 222 and 4,004 Hz; 12% steps). REG patterns consisted of 20 frequencies randomly selected from the pool and iterated over 3 to 4 cycles. In the 5^th^ block, each REGr was randomly transposed up or down by one tone (12%; shifted 1 place higher or lower in the frequency pool than the original, see Fig. S6C). To allow for the transposition, REGr patterns were drawn from a subset of 24 frequencies (i.e., not including the highest and lowest frequency in the pool).

##### Participants

Twenty paid individuals (twelve females; average age, 24.75 ± 6.8 years) took part in the study. No participant reported hearing difficulties.

#### Results

Overall, the same pattern of performance as in Exp. 1A was demonstrated. Fig. S6B demonstrates progressively stronger implicit memory for REGr, as revealed by a growing RT advantage over novel REG across blocks [main effect of condition: F(1,19) = 47.31, p < .001, η_p_^2^ = .71; main effect of block: F(4, 76) = 7.95, p < .001, η_p_^2^ = .29; interaction condition per block: F(4, 76) = 5.35, p = .003, η_p_^2^ = .22]. Specifically, whilst in the first block performance didn’t differ between RANREG and RANREGr conditions [t(19) = 1.635, p = .59], a significantly faster response (186 ms; 3.7 tones) for RANREGr was observed in the second block [t(19) = 4.302 p = .001], and grew across the remaining blocks (all ps < .004).

Importantly, this RT advantage (205 ms – 4.1 tones) in block 5 (transposed REGr) didn’t differ from the RT advantage on block 4 (272 ms; 5.4 tones) [t(19) = 1.541, p = .14]. To confirm the immediacy of the transfer we compared the RT advantage in the first intra-block presentation in block 5, where the transposition was introduced, with the third (last) intra-block presentation in block 4 (Fig S1H). No difference was observed [t(19) = 1.26, p =.223; Fig. S1H], suggesting that the generalization to the transposed pattern was instantaneous.

The observation of a transfer of RT advantage to the transposed sequences may suggest that the formed representation is not precisely echoic: instead of the specific frequency pattern, the auditory system might be maintaining a representation of the contour, or inter-tone interval within the REGr pattern. Another possibility is that the tolerance reflects a noisy frequency representation, though we note that the frequency steps here (12%) are large enough to be discriminable by most listeners.

**Fig. S6.**
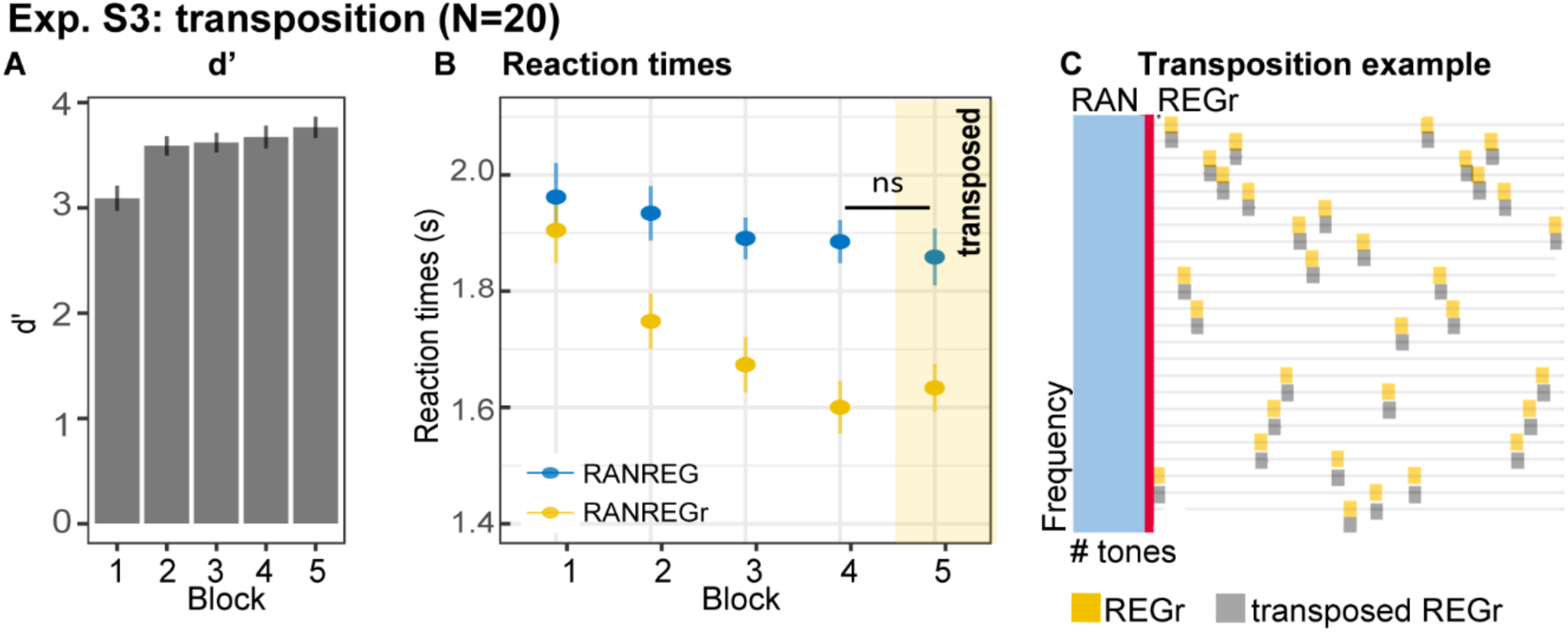
Experiment S3: implicit memory is robust to pattern transposition. **(A)** d’ across all blocks. Error bars indicate 1 s.e.m. **(B)** RT to the transition in RANREG and RANREGr across blocks. In block 5 (yellow shading) the originally learned REGr were replaced by transposed versions. Error bars indicate 1 s.e.m. **(C)** Schematic example of the transposition. Yellow squares indicate tones in a REGr sequence; grey squares indicate the transposed version (in this example, the REG tones were shifted downwards by one step in the frequency pool; 12%). The red line indicates the transition from RAN (light blue area) to REGr

## Notes

https://osf.io/dtzs3/

